# Effects of Perturbation Velocity, Direction, Background muscle activation, and Task Instruction on Long-Latency Responses Measured from Forearm Muscles

**DOI:** 10.1101/2020.10.12.335869

**Authors:** Jacob Weinman, Paria Arfa Fatollahkhani, Andrea Zonnino, Rebecca Nikonowicz, Fabrizio Sergi

## Abstract

The centeral nervous system uses feedback processes that occur at multiple time scales to control interactions with the environment. Insight on the neuromechanical mechanisms subserving the faster feedback processes can be gained by applying rapid mechanical perturbations to the limb, and observing the ensuing muscle responses using electromyography (EMG). The long-latency response (LLR) is the fastest process that directly involve cortical areas, with a motorneuron response measurable 50 ms following an imposed limb displacement. Several behavioral factors concerning perturbation mechanics and the active role of muscles prior or during the perturbation can modulate the long-latency response amplitude (LLRa) in the upper limbs, but the interaction between many of these factors had not been systematically studied before.

We conducted a behavioral study on thirteen healthy individuals to determine the effect and interaction of four behavioral factors -- background muscle torque, perturbation direction, perturbation velocity, and task instruction -- on the LLRa evoked from the flexor carpi radialis (FCR) and extensor carpi ulnaris (ECU) muscles following the application of wrist displacements. The effects of the four factors listed above were quantified using both a 0D statistical analysis on the average perturbation-evoked EMG signal in the period corresponding to an LLR, and using a timeseries analysis of EMG signals.

All factors significantly modulated LLRa, and that their combination nonlinearly contributed to modulating the LLRa. Specifically, all the three-way interaction terms that could be computed without including the interaction between instruction and velocity significantly modulated the LLR. Analysis of the three-way interaction terms of the 0D model indicated that for the ECU muscle, the LLRa evoked when subjects are asked to maintain their muscle activation in response to the perturbations (DNI) was greater than the one observed when subjects yielded (Y) to the perturbations (ΔLLRa — DNI vs. Y: 1.76±0.16 nu, p<0.001), but this effect was not measured for muscles undergoing shortening or in absence of background muscle activation. Moreover, higher perturbation velocity increased the LLRa evoked from the stretched muscle in presence of a background torque (ΔLLRa 200−125 deg/s: 0.94±0.20 nu, p<0.001; ΔLLRa 125−50 deg/s: 1.09 ±0.20 nu, p<0.001), but no effects of velocity were measured in absence of background torque, nor effects of any of those factors was measured on muscles shortened by the perturbations. The time-series analysis indicated the significance of some effects in the LLR region also for muscles undergoing shortening. As an example, the interaction between torque and instruction was significant also for the ECU muscle undergoing shortening, in part due to the composition of a positive and negative modulation of the response due to the interaction between of the two terms. The absence of a nonlinear interaction between task instruction and perturbation velocity suggest that the modulation introduced by these two factors are processed by distinct neural pathways.

## Introduction

Countering unexpected and unpredictable loads is a ubiquitous occurrence of everyday life. Humans can precisely perform movements and interact with the environment even in the presence of these external perturbations. These mechanical perturbations require the nervous system to induce a compensatory action in order to ensure the task success. An important component of the compensatory actions produced by the central nervous system is the long-latency response (LLR). In upper limb muscles, the LLR is evident as the burst of muscle activity occurring 50–100 ms following a limb displacement. Accordingly, this event occurs between the fastest nervous system response, i.e., the short-latency reflex (SLR) occurring within 20-50ms, and the delayed voluntary reaction which begins 100 ms after the imposed perturbation (1–3).

After a seminal study by Hammond in 1956 (1), several investigators have utilized a limb perturbation paradigm to investigate the physiological mechanisms subserving the muscle stretch responses to the externally applied loads (3–8). In these paradigms, the muscle responses including the LLRs are recorded through surface electromyography (EMG) activity evoked in the muscle stretched by an imposed angular joint displacement induced by a mechanical perturbation of known and controllable characteristics (3,9–14).

Several behavioral factors are known to affect the amplitude of LLRs, including the neuromechanical state of the muscle prior to the application of a perturbation (i.e., muscle length and activation) (15,16), the direction of perturbation (i.e., whether the perturbation stretches or shortens the muscle) (17–19), the kinematic features of the applied perturbation (i.e., perturbation velocity, duration, amplitude, velocity profile) (20–22), and the instructions provided to participants as to how to respond to the applied perturbations (19,23,24).

Although investigators mostly focused on studying LLR features in stretched muscles, there is evidence of EMG activity evoked in the muscle shortened by the applied perturbation (18,19). Specifically, an increase in the EMG activity of the extensor carpi radialis (ECR) muscle which was shortened due to the applied wrist extension perturbation was documented in a study conducted in 2004 (18). Although the shortened muscle response was smaller in amplitude than the one evoked in the stretched (flexor) muscle, both had two separate components of SLR and LLR with a very similar onset timing. Another study also observed a similarly timed low-amplitude EMG response with an onset of about 50ms, evoked in the ECR muscle in response to a rapid wrist extension displacement (19). The authors suggest that part of the measured effects may be due to cross-talk (a volume-conducted response from the stretched muscle). However, because significant perturbation-evoked responses were measured in muscles undergoing shortening even using intramuscular EMG (17), it may be reasonable that LLR are evoked in muscles subject to both a shortening and a stretching perturbation.

Studies also examined the effects of background muscle activation prior to the imposed perturbation on the LLR amplitude. In general, an increase in background activation results in an increase in the magnitude of muscle activity in both the proximal and distal muscles of the upper limbs (15,19,25). Affecting the background motoneuron pool excitability, pre-existing background muscle activation is thought to reflect an automatic adjustment mechanism, known as the automatic gain component of the LLR (15,19,25,26).

Several studies quantified the effects of the kinematic features of the applied perturbation on the LLR amplitude (2,20–23). It is generally accepted that the LLR amplitude increases as a function of the velocity of the applied perturbations (15,27). In the common ramp-and-hold perturbation paradigms, which are conducted at constant velocity, perturbation duration may also play a factor when the duration of the perturbation is within the range of neuromuscular delays expected for the LLR. However, the details of the interaction between perturbation velocity and duration in modulating LLR amplitude are not yet completely understood. A study by Lewis et al. showed that LLR amplitudes of the biceps brachii undergoing stretch are modulated by velocity in all conditions, but the slope of the relationship is also modulated by duration (21). Yet, the range of velocities and durations that modulate the response in such a way is likely limited. In fact, we know that for FCR, a very high velocity and short duration (<40ms) perturbation is not sufficient to evoke a long-latency response, whereas a perturbation of low velocity and long duration (>60ms) generates well-developed LLRs (2).

Task instruction also plays a key role in modulation of the LLR response. Previous studies showed that larger LLRs are evoked when participants attempt to counter a perturbation than yield to the perturbation (16,19,23,24). Participants can be instructed to respond to the perturbation in different ways: they can be asked to relax immediately following the perturbation (19) --- a condition referred to as “yield”; to maintain the background torque and avoid a voluntary response to the perturbation (16) --- a condition referred to as “Do Not Intervene”; or to explicitly compensate by activating their muscles in the opposite direction of the perturbation (23), or to compensate cued by a visual feedback of the hand position (24) --- conditions referred to as “Resist”. The experiments where the subject was instructed to either counter the stretch perturbation or yield to it demonstrated that the LLR can be modulated to functionally adapt to the task at the upper limb (5,13,28). Studies also suggest that the LLR is subject to an internal model of the limb configuration, which increases the functional effectiveness of the muscle response to the external loads (29). Moreover, accumulating evidence shows that the temporal overlap of two different responses including a task-dependent response and an automatic response results in the task-dependent change in LLR amplitude (13,23,30).

Gathered together, several factors concerning the mechanics of the applied perturbations and the active role of muscles prior or during the perturbation can modulate the amplitude of long-latency responses in the upper limbs. Hence, it is of a paramount importance to study how the interaction of these factors would affect the muscle stretch responses in a single study. However, the majority of previous studies has systematically studied only one or two of the factors modulating LLRa, with the interaction between perturbation velocity and task instruction studied in (23), the interaction between perturbation velocity and background torque studied in (15), and the interaction between task instruction and background torque studied in (16). One previous study (19) has studied the effects of three of the factors highlighted above (i.e., task instructions, perturbation direction, and torque), though not with a full factorial design capable of quantifying the interactions between all factors. As such, to the best of our knowledge, no previous study conducted a full factorial design capable of quantifying the effects of and the interactions between task instructions, perturbation duration, background torque, and perturbation velocity on LLR amplitude.

There is currently limited knowledge on the neural substrates of LLR, and specifically on the contributions of subcortical areas to the generation of LLRs (3). A possible method to study subcortical contributions to LLRs is to combine robotic perturbations with surface EMG and fMRI, an approach that our group has recently started to pursue (8). Our proposed method is based on perturbations delivered to the wrist because the muscles are farther away from the scanner isocenter during fMRI, and because wrist movements minimally induce head movements and distort the magnetic field in the imaging region. However, to design an experiment where stretch-evoked responses are measured in a way that the ensuing neural responses are maximally decoupled and thus identifiable via fMRI, it is important to first establish the effects of relevant behavioral conditions on the expected LLR amplitude. Unfortunately, the effects of some of the behavioral factors reported above have not been investigated for the wrist joint, such as the interaction between perturbation direction and velocity.

The goal of this study is to determine the effects of and the interactions between several experimental factors modulating the LLR amplitude during ramp-and-hold perturbation. Specifically, the goal of this study is to establish the effect and interaction of background muscle torque, perturbation direction, perturbation velocity, and task instruction on the LLR amplitude evoked from the flexor carpi radialis (FCR) and extensor carpi ulnaris (ECU) muscles following the application of controlled angular displacements of the wrist in both extension and flexion direction.

## Materials and Methods

Thirteen healthy individuals were recruited to participate in this study (protocol approved by the University of Delaware Institutional Review Board, protocol no. 1097082-6). Subjects – age (mean ± s.d.: 24±3 years) were naïve to the purpose of the study and free from known neurological or orthopedic disorders affecting arm function. Subjects were exposed to an experiment that aimed to quantify the amplitude of long-latency responses via the recording of EMG activity from a wrist flexor and extensor pair, the flexor carpi radialis (FCR) and extensor carpi ulnaris (ECU). These responses were evoked flexion or extension perturbations applied by a robot to a subject’s wrist in various conditions.

### Materials

#### Perturbation robot

A custom-developed robot, the MR-StretchWrist, was used to apply perturbations to subject’s wrists. The MR-StretchWrist is a 1-degree of freedom robot that can provide wrist flexion and extension between −45 to 45 degrees (8). The robot employs an ultrasonic piezoelectric motor (EN6060, Shinsei Motor Inc., Japan) that can provide 500 mN·m of torque and can move at velocities of up to 900 degrees/second.

To provide torque for sufficient muscle stretch within the desired time of 50 ms, a capstan transmission with a 3:1 gear ratio was included in the design. The capstan drive contains two pulleys with different diameters connected via a microfiber braided line (SpiderWire Stealth SPW-0039, 0.4mm diameter braided fishing line). The cable is wrapped around each pulley multiple times to ensure zero slippage. The capstan transmission is an ideal candidate for this application because it has no backlash, low friction, and high bandwidth.

Measurement of the wrist flexion/extension angle was obtained using an incremental encoder (resolution: 0.09 deg) placed on the motor shaft, with a resulting resolution in measuring the wrist flexion/extension angle of 0.03 deg.

#### EMG amplifier and electrodes

EMG data was recorded with an OTBioelettronica EMG-USB2+ amplifier (OTBioelettronica s.r.l., Torino, Italy), using OTBiolab Software (OTBioelettronica s.r.l., Torino, Italy). Disposable Silver/Silver chloride surface electrodes with conductive gel (HEX Dual Electrodes, Noraxon USA, Scottsdale, AZ) were placed on the skin of the subject. A moistened conductive band was wrapped around the wrist, ensuring contact with the radial and ulnar stylar processes, to serve as a reference electrode. Bipolar cables were attached to the disposable electrodes and connected to the amplifier.

#### Force measurement

An ATI Mini 27 Ti force sensor integrated in the MR-StretchWrist is used to measure the torque applied by the subject in the z direction (Full-scale Load (FSL): 2 Nm, Resolution: 0.5 mNm, Measurement uncertainty: 1.5% of FSO). Transducer signals are processed by a preamplifier box (ATI Industrial Automation, Apex, NC), and digitized by an analog/digital I/O device (PCIe-6321, National Instruments, Austin, TX) connected to a standard desktop computer.

#### Control software

Software for robot position control, perturbation timing, and for the graphical user interface was developed in Matlab and Simulink, and executed in real-time (sample rate: 1 kHz) using the QUARC real-time control software (Quanser, Markham, ON). Encoder data were acquired using the Q2 USB data acquisition board (DAQ). The experimental setup is shown in Fig. 1.

**Fig. 1:**
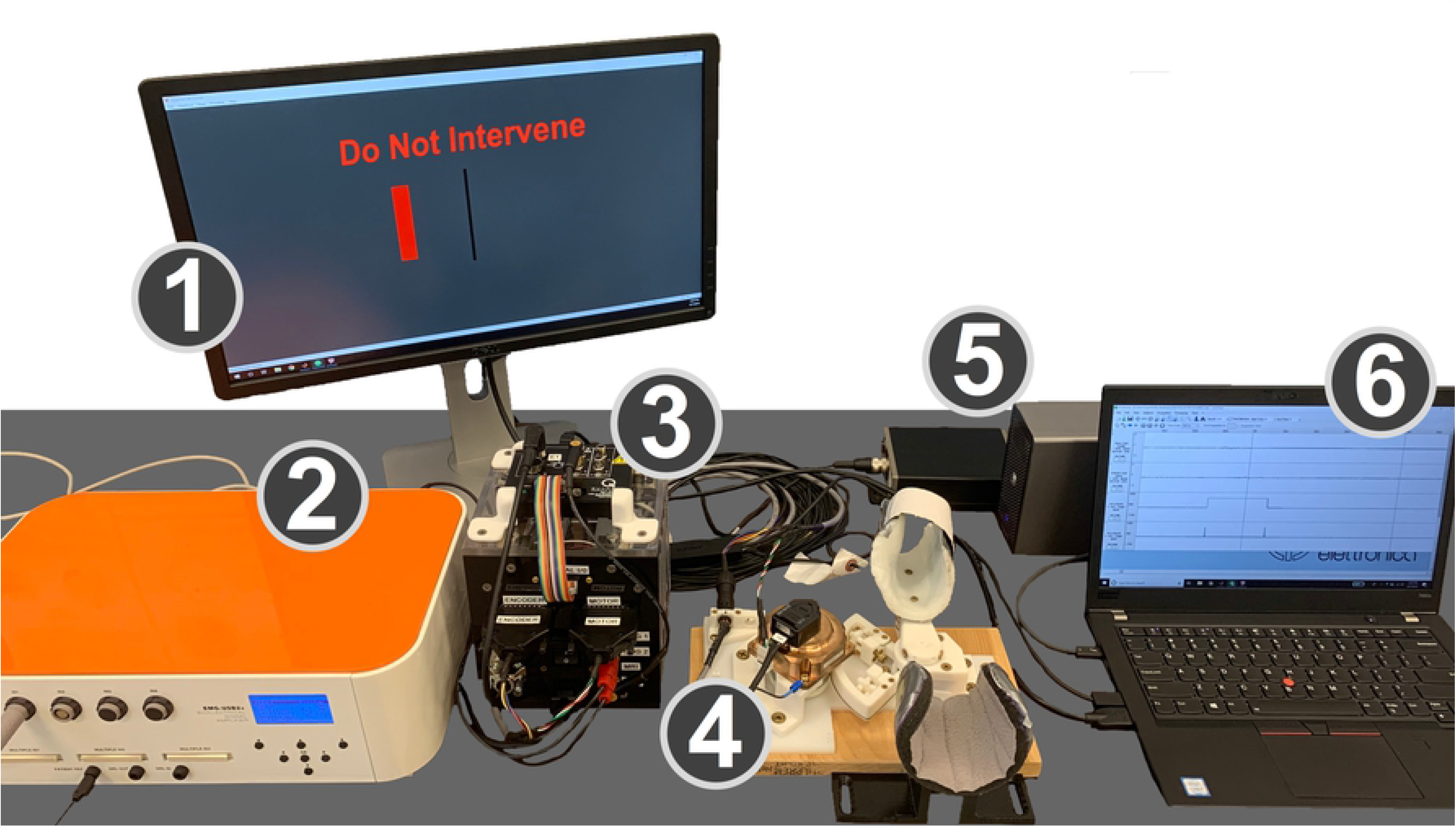
Experimental setup. (1) Monitor displaying the GUI that cued subjects to the desired level of wrist flexion or extension torque and provides task instructions. (2) EMG amplifier. (3) Control box with the power supply, motor driver, and data acquisition board. (4) MR-StretchWrist robot. (5) Force sensor preamplifier box and analog/digital I/O device for force sensor data. (6) Laptop running EMG collection software, Simulink, and real-time QUARC software.

### Methods

#### EMG electrode positioning

To determine the location of the electrodes to measure activity of the FCR and ECU, manual palpations were performed on the right arm of the subjects. Repeated wrist flexion during palpation while the wrist was in its neutral position aided in locating the FCR. Similarly, repeated wrist extension in this same orientation aided in locating the ECU. Points parallel to the muscle’s fibers were drawn 3 cm apart. The skin was then prepped with 70% isopropyl alcohol wipes and the application a thin layer of conductive skin prep gel (Nuprep, Weaver and Co., Aurora, CO).

#### Experimental procedures

In this study, subjects were seated with their forearms resting in a stationary support connected to a normal desk, with the forearm extending anteriorly in front of the body. Their hand was strapped inside of a mold such that their wrist was in a semiprone/neutral condition, and any static wrist flexion/extension torque (range: 0 – 2 Nm) would be supported by the mold with little deflections. A GUI was shown on a computer screen indicating the amount of wrist flexion/extension torque to apply. After the subject reached the appropriate torque target, and maintained it within a 50 mNm range for 0.4 to 0.8 seconds, the robot perturbed the wrist in the direction that would stretch the agonist muscle with respect to the cued torque (i.e., if the background torque was flexion, the perturbation was wrist extension). Perturbations were applied for a duration of 200 ms in all conditions, so to avoid undesired oscillations of the EMG signal due to impact dynamics arising from the abrupt end of a perturbation. After each wrist perturbation, the robot halted for 1 second before returning to the neutral position for the following perturbation. Numerous conditions were studied in this protocol, defined by factorial combinations of four factors: 1) perturbation velocity, 2) perturbation direction, 3) background torque, 4) task instructions.

Perturbation velocity assumed three levels: 50, 125, and 200 deg/s. Perturbation direction assumed two levels: wrist flexion or wrist extension. Background torque was set to either 0 or 200 mNm. Task instruction assumed two levels: “yield” (Y) and “do not intervene” (DNI). In the Y condition, subjects were told to not provide any resistance after the perturbation and yield (i.e., relax) to the movement. In the DNI condition, subjects were told to continue applying the same amount of torque that they were applying prior to perturbation. These two instructions are fundamentally the same in the absence of a background torque.

Ten trials per condition (velocity, direction, background torque, instruction) were collected, resulting in each experiment consisting of a total of 240 perturbations. The order in which conditions were applied was randomly generated by the Simulink and MATLAB files. Furthermore, the time between the end of the prior perturbation and the start of the next perturbation cue was randomized between 3 and 7 seconds.

#### EMG processing

Pre-processing of the raw EMG data was conducted using Matlab code (Matlab 2017a, MathWorks, Natick, MA). A band-pass filter, a 4th order Butterworth filter (f_LP_=250 Hz and f_HP_=20 Hz), was used to remove low frequency noise related to movement artifacts and high frequency noise related to intrinsic measurement noise. Matlab’s *filtfilt* function was used to guarantee a filtered waveform without any phase shift of the signal. The filtered waveform was then rectified. Digital outputs produced by the DAQ board (Q2-USB, Quanser, Markham, ON) were sent to the EMG amplifier to identify instants of perturbation onset, used for segmentation of the EMG signal into 200 ms long timeseries, one for each perturbation, each starting at the time of perturbation onset. EMG tracks were shifted in time such that t=0 ms corresponded to the instant of arrival of the pulse.

The amplitude of the segmented EMG signal measured during a perturbation was normalized by the average magnitude of the rectified EMG signal measured from that same muscle during agonist background contractions preceding all perturbations. This procedure allowed us to conduct group analysis of normalized EMG data. With this procedure, an EMG signal of unitary magnitude indicated that the rectified EMG signal measured during each perturbation had the same amplitude of the average EMG signal generated by that muscle for a 200 mNm isometric torque. Units of the normalized EMG timeseries are referred to as normalized units (nu).

These procedures resulted in 10 segmented timeseries extracted from each subject per combination of conditions (velocity, direction, torque, instruction). Long-latency response amplitude (LLRa), defined in agreement with the literature (31) as the average signal of the EMG tracks in the [50 100] ms interval, was calculated for each perturbation for both muscles, and indexed as a function of subject, repetition, and combination of experimental conditions. The subject-specific average of LLRa for each combination of perturbation conditions was used as the outcome measure for the 0-D statistical analysis.

Repeated measurements from each subject were averaged to yield the timeseries with average rectified EMG response for each subject EMG_*sub,v,d,t,inst*_ used for the 1-D statistical analysis. Group averages 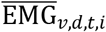 and corresponding standard deviations 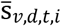 were then calculated for display purposes.

#### Statistical analysis

Two four-way full factorial linear mixed model ANOVAs were conducted using the subject-specific average LLRa measured for FCR and ECU, respectively, as outcome measure. The four factors included in the ANOVA were perturbation velocity (0, 125, 200 deg/s), background torque (0, 200 mNm), instruction (yield, do not intervene), and perturbation direction (stretch vs. shorten), defined based on the effect that the perturbation would have on the length of each muscle. As an example, a flexion perturbation would correspond to the “stretch” level for ECU, and the “shorten” level for FCR. Statistical analysis was conducted using JMP, and all variables were coded as nominal variables, with the default conditions of 50 deg/s for velocity, 0 mNm for background torque, yield for instruction, non-stretch for perturbation direction. JMP Pro Version 14 (SAS Institute Inc., Cary, NC, USA) was used for this analysis.

The linear mixed model included 16 terms for fixed effects (4 main effects, 6 two-way interactions, 4 three-way interactions, one four-way interaction term, and an intercept), plus an additional set of offset variables for random subject-specific effects. The Satterthwaite method was used to determine the number of degrees of freedom in the model. All terms are reported if their estimated effect is significant at the type I error rate α=0.05. In those cases, Tukey HSD post-hoc tests were also used to determine pairs of levels with significant differences.

A 1-D ANOVA was also conducted on the timeseries of rectified EMG signal (EMG_*sub,v,d,t,i*_) measured during the post-perturbation interval comprised between 0 and 200 ms using spm1d software. 1-D statistical analysis models are useful in this application to analyze the effects of the experimental conditions on the perturbation-induced muscle response without prior hypotheses on the specific time interval where an effect is expected (32,33). While with the 0D analysis we restricted our focus on the time interval ensuing the perturbation comprised between 50 and 100 ms (thus obtaining a scalar, or 0D, outcome measure), with the 1D model we sought to determine whether there is an effect on the timeseries of measured EMG amplitudes associated with all the experimental conditions, and their combinations, at *any* time point in the post-perturbation interval comprised between 0 and 200 ms.

Because the current version of the spm1D Matlab software only allows to build full factorial models with a maximum of three main effects, we broke each four-way model into two three-way models and performed our analysis using the spm1d function *anova3rm*. Each model included the factors speed, background torque, and instruction, with one model including LLRs measured during stretch, and the other model including LLRs measured during shortening. The spm1d analysis performs a mixed model ANOVA for measurements collected at all time points, and then conducts inference controlling for the associated multiple corrections using random field theory (32). As such, the output of the 1D ANOVA procedure is a time-series of F scores for all main effects and their interaction, combined with the identification of time intervals where those effects are significant at α=0.05. When an effect or an interaction was significant during the time-window where an LLR is expected (i.e., from 50 to 100 ms after perturbation onset), we conducted post-hoc tests to determine pairs of levels with significant modulation in EMG signal.

## Results

Of the 13 participants recruited for this study, data sets of 2 subjects for ECU and 3 subjects for FCR were excluded from analyses due to poor EMG recordings. Therefore, the 0-D analysis and the 1D analysis were performed on data collected on *n=11* individuals for the ECU muscle, and *n=10* individuals for the FCR muscle. Timeseries extracted in the different experimental conditions for the ECU muscle are shown in Fig. 2 and Fig. 3. Similar representations are provided for the FCR muscle in Fig. S1 and Fig. S2 of the Supplementary Materials.

**Fig. 2.**
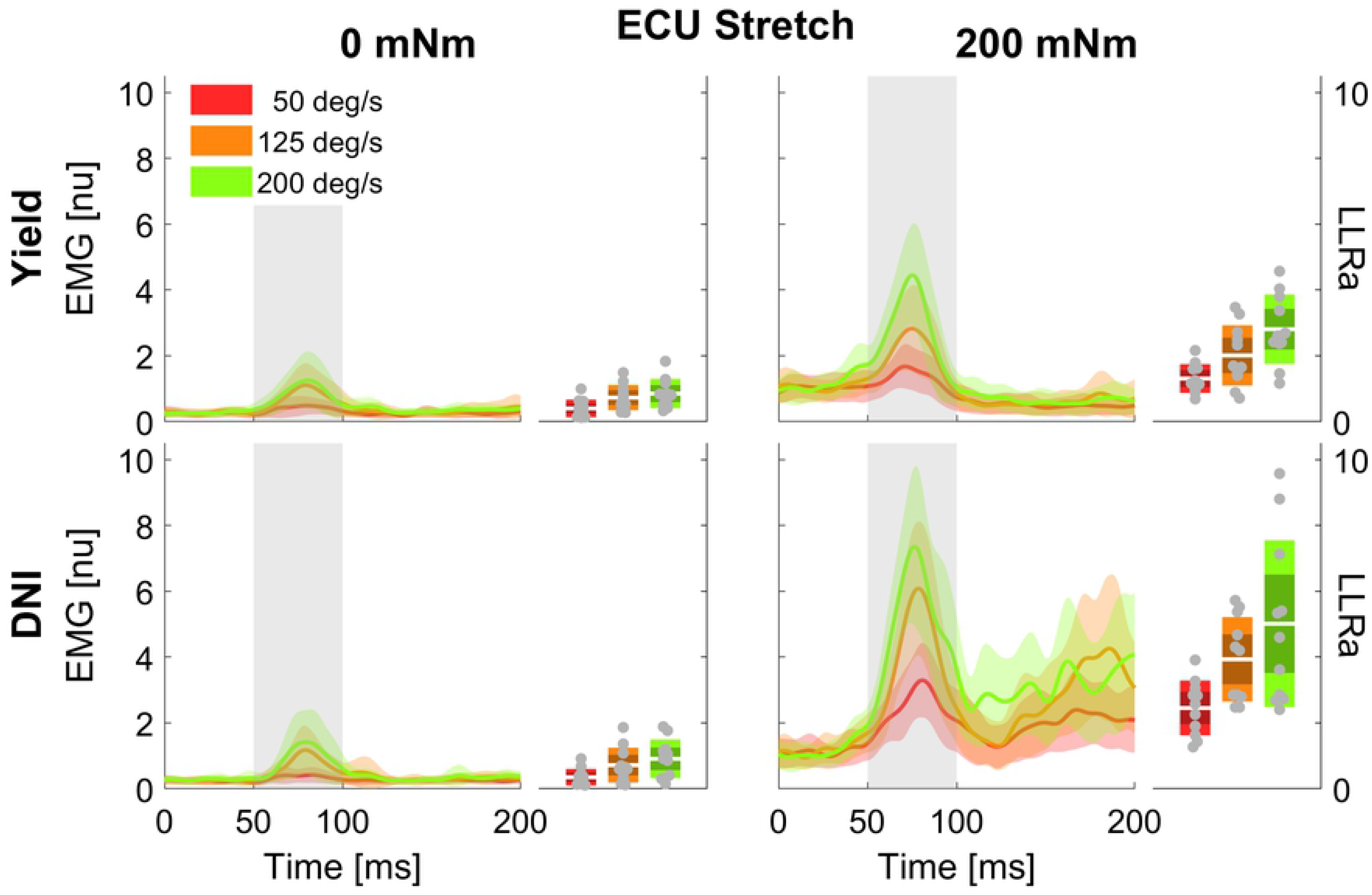
Time series of EMG signal (left half-panels) and LLRa (boxplot in the right half-panels) from all subjects resulting with perturbations in the flexion direction (stretching the ECU muscle). Color indicates the speed of the perturbation (red = 50 deg/s, orange = 125 deg/s, green = 200 deg/s), with the line indicating the mean and the shaded area indicating the ±1 s.d. region. Panels in different rows include measurements at two levels of task instruction – Yield (top), and DNI (bottom); columns include measurements at two levels of background torque – 200 mNm (right), and 0 mNm (left). In the boxplots, darker shades represent 1 standard deviation from the mean, and lighter shades indicate the 95% confidence interval

**Fig. 3.**
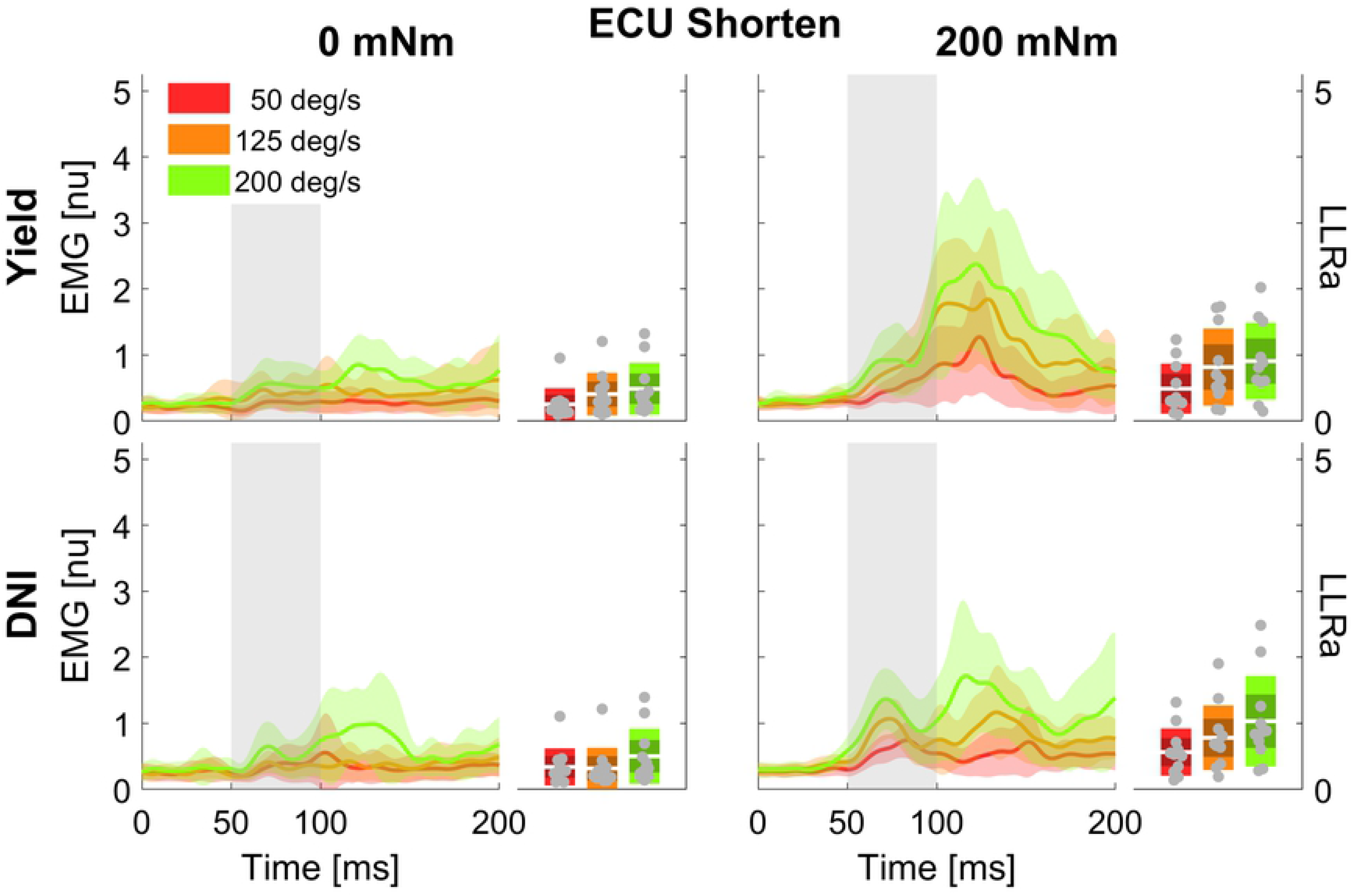
Time series of EMG signal and LLRa from all subjects resulting with perturbations in the extension direction (shortening the ECU muscle).

### 0-D analysis

The linear mixed model computed an adjusted R^2^ of 0.627 for FCR, and an adjusted R^2^ of 0.788 for ECU. The model reported a significant effect of all four main factors. However, since all factors are involved in several two- and three-way interactions, only the interactions will be analyzed and discussed below. Results are presented below as least square means ± standard error (LLRa) or difference in least square means ± standard error (ΔLLRa) in units of the outcome measure, i.e. normalized units (nu). A report including the significant fixed effects from the linear mixed model ANOVA is provided below in Table 1 and Table 2.

**Table 1.**
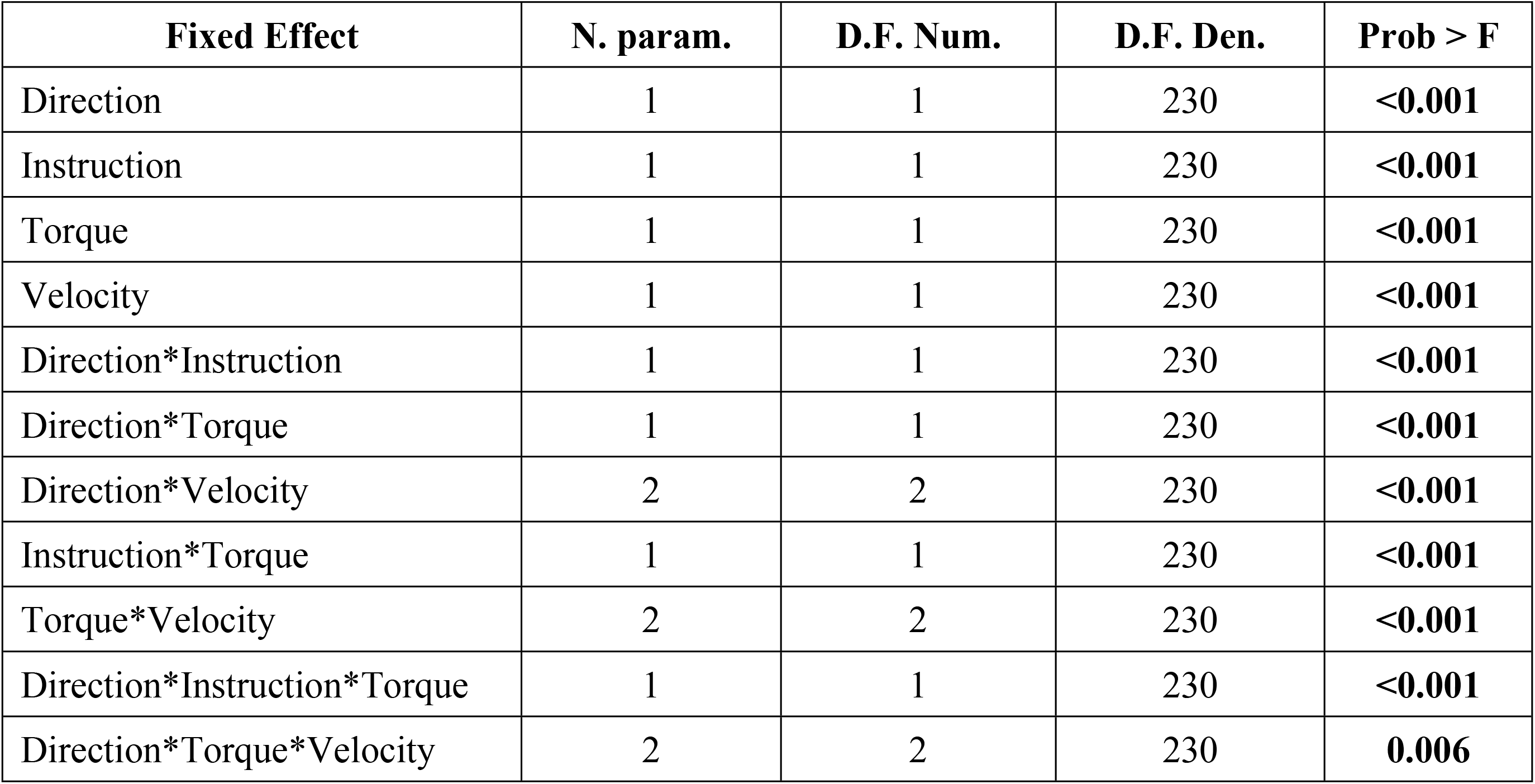
Significant main effects and interactions for ECU LLRa. A total of 11 effects are presented.

| **Fixed Effect** | **N. param.** | **D.F. Num.** | **D.F. Den.** | **Prob > F** |
| --- | --- | --- | --- | --- |
| Direction | 1 | 1 | 230 | **<0.001** |
| Instruction | 1 | 1 | 230 | **<0.001** |
| Torque | 1 | 1 | 230 | **<0.001** |
| Velocity | 1 | 1 | 230 | **<0.001** |
| Direction*Instruction | 1 | 1 | 230 | **<0.001** |
| Direction*Torque | 1 | 1 | 230 | **<0.001** |
| Direction*Velocity | 2 | 2 | 230 | **<0.001** |
| Instruction*Torque | 1 | 1 | 230 | **<0.001** |
| Torque*Velocity | 2 | 2 | 230 | **<0.001** |
| Direction*Instruction*Torque | 1 | 1 | 230 | **<0.001** |
| Direction*Torque*Velocity | 2 | 2 | 230 | **0.006** |

**Table 2.**
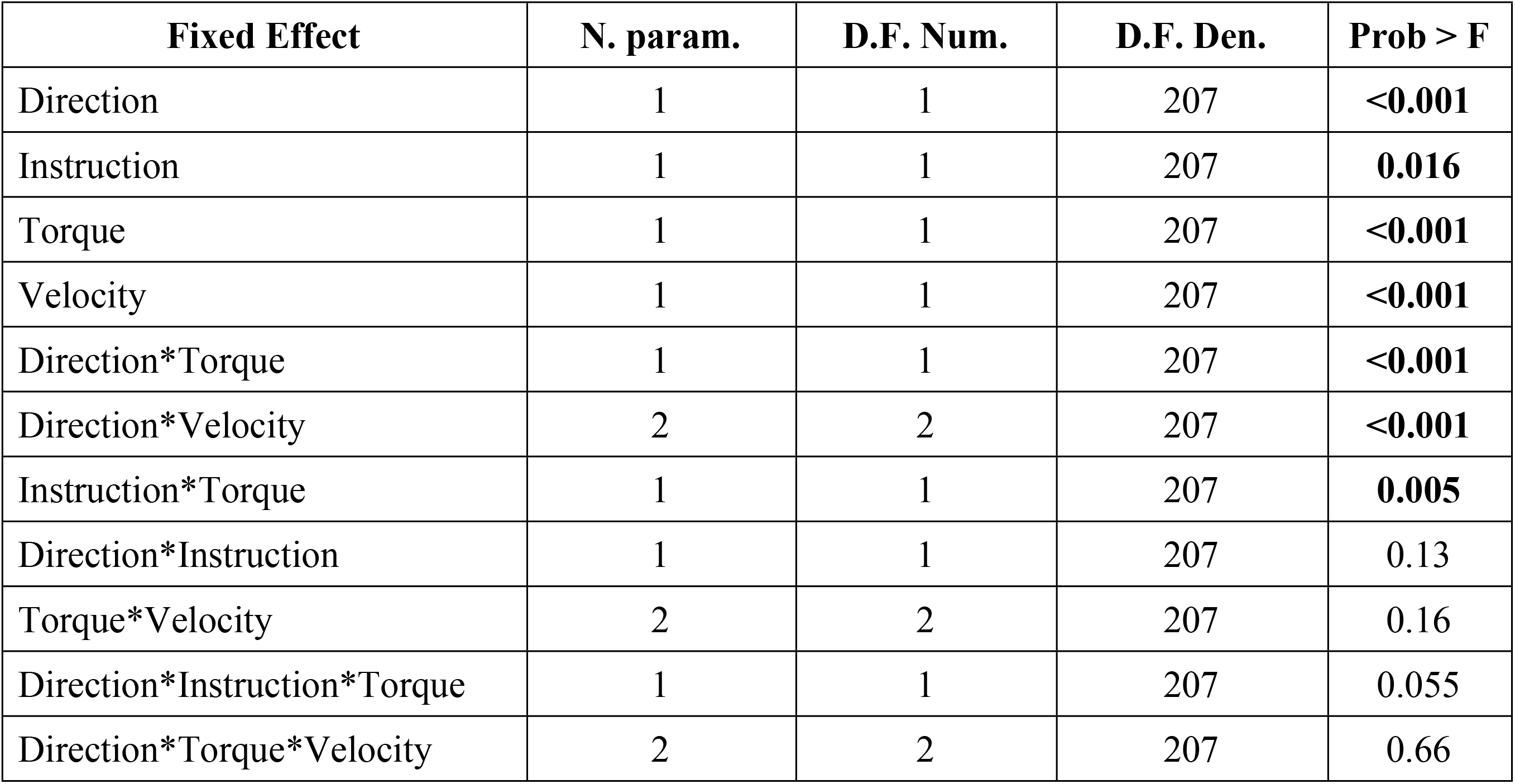
Main effects and interactions for FCR LLRa. Of the 11 effects listed from Table 1, the FCR LLRa shares 7 significant effects.

| **Fixed Effect** | **N. param.** | **D.F. Num.** | **D.F. Den.** | **Prob > F** |
| --- | --- | --- | --- | --- |
| Direction | 1 | 1 | 207 | **<0.001** |
| Instruction | 1 | 1 | 207 | **0.016** |
| Torque | 1 | 1 | 207 | **<0.001** |
| Velocity | 1 | 1 | 207 | **<0.001** |
| Direction*Torque | 1 | 1 | 207 | **<0.001** |
| Direction*Velocity | 2 | 2 | 207 | **<0.001** |
| Instruction*Torque | 1 | 1 | 207 | **0.005** |
| Direction*Instruction | 1 | 1 | 207 | 0.13 |
| Torque*Velocity | 2 | 2 | 207 | 0.16 |
| Direction*Instruction*Torque | 1 | 1 | 207 | 0.055 |
| Direction*Torque*Velocity | 2 | 2 | 207 | 0.66 |

A significant three-way interaction between perturbation direction, instruction, and torque was measured for the ECU muscle (Fig. 4). A significant effect of instruction or perturbation direction on LLRa was observed only in the presence of a background torque for both muscles. In presence of background torque, the LLRa associated with stretch perturbations was greater than the LLRa associated with shortening perturbation both when task instructions were DNI (LLRa — Stretch DNI: 3.80±0.17 nu, Shorten DNI: 0.79±0.17 nu, p<0.001), and when task instructions were to yield (LLRa — Stretch Y: 2.04±0.17 nu, Shorten Y: 0.74±0.17 nu, p<0.001). In presence of background torque and muscle stretch, LLRa was greater in the DNI condition compared to yield (ΔLLRa — DNI vs. Y: 1.76±0.16 nu, p<0.001).

**Fig. 4.**
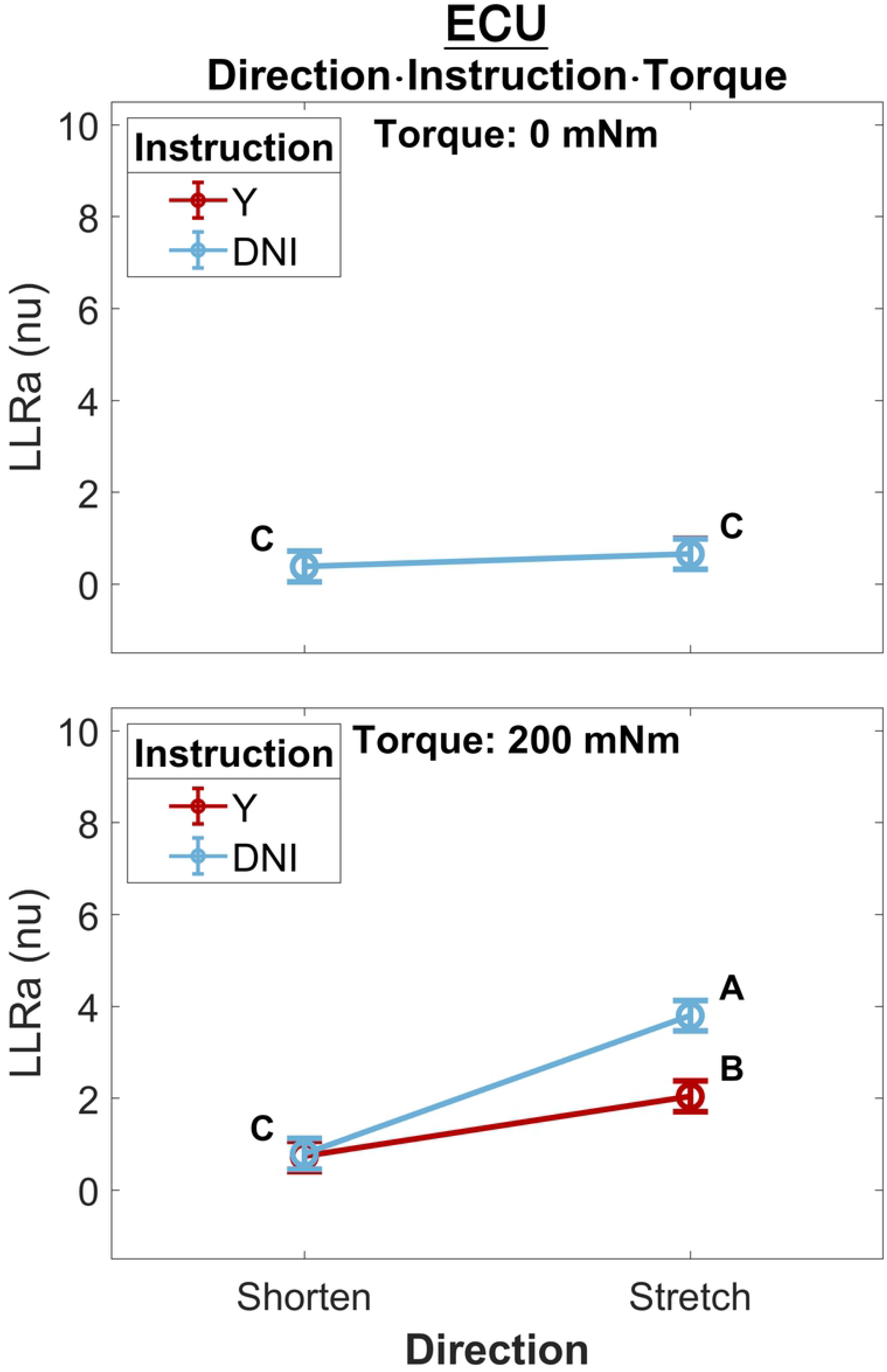
Least square means (mean displayed with a circle and whisker extending to ± one standard error of the mean) for the significant three-way interaction between perturbation direction, instruction, and torque, conducted for the ECU muscle. Plots are split by torque level -- 0 mNm (top), 200 mNm (bottom). Line colors indicate instruction, and different levels of perturbation direction are displayed along the x axis. Letters on the plots are representative of post-hoc Tukey HSD tests; pairs of elements that do not share a letter are significantly different.

A significant three-way interaction between perturbation direction, torque, and velocity was measured for the ECU muscle (Fig. 5). A significant effect of perturbation velocity or direction was observed only in the presence of a background torque. In presence of background torque, the LLRa associated with stretch perturbations was greater than the LLRa associated with shortening perturbations at all velocities (200 deg/s: Stretch: 3.91±0.19 nu, Shorten: 0.97±0.19 nu, p<0.001; 125 deg/s: Stretch: 2.97±0.19 nu, Shorten: 0.80±0.19 nu, p<0.001; 50 deg/s: Stretch: 1.88±0.19 nu, Shorten: 0.53±0.19 nu, p<0.001). In presence of background torque and for perturbations that stretched the muscle, LLRa increased with velocity (ΔLLRa 200−125 deg/s: 0.94±0.20 nu, p<0.001; ΔLLRa 125−50 deg/s: 1.09 ±0.20 nu, p<0.001). Instead, no velocity-dependent effect was measured for perturbations shortening the muscle.

**Fig. 5.**
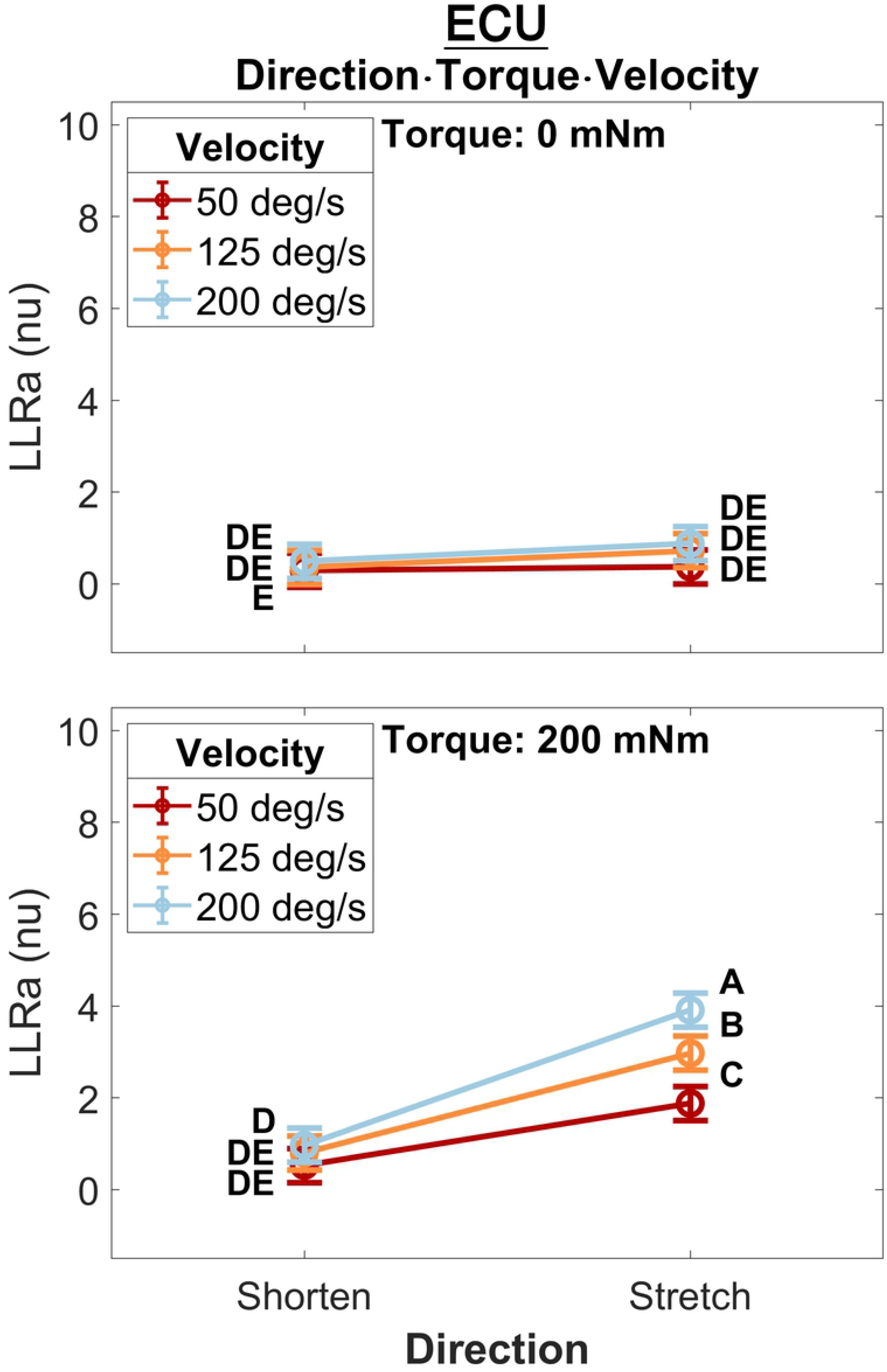
Least square means for the significant three-way interaction between perturbation direction, velocity, and torque, conducted for the ECU. Plots are split by torque level -- 0 mNm (top), 200 mNm (bottom) -- line color indicates velocity, and different levels of perturbation direction are displayed along the x axis.

Five two-way interaction terms were significant for ECU, while three terms were significant for FCR. Three terms were common to both muscles and are described next.

The two-way interaction between instruction and torque resulted from a greater increase in LLRa measured in the DNI conditions compared to the yield conditions measured in the presence of a background torque for both muscles (Fig. 6). In the presence of background torque, LLRa measured during the DNI condition was significantly greater than LLRa measured in the Y condition (FCR LLRa — DNI: 3.67±0.53 nu, Y: 2.39±0.53 nu, p=0.001; ECU LLRa — DNI: 2.30±0.15 nu, Y: 1.40±0.15 nu, p<0.001). There was no difference in LLRa measured in absence of background torque between the two instructions (FCR LLRa — DNI: 1.12±0.53 nu, Y: 1.22±0.53 nu, p=0.990; ECU LLRa — DNI: 0.52±0.15 nu, Y: 0.52±0.15 nu, p=1). Significant differences between torque levels were measured in both the FCR and ECU for both instructions (FCR ΔLLRa — DNI: 2.55±0.34 nu, p<0.001, Y: 1.16±0.34 nu, p=0.004; ECU ΔLLRa — DNI: 1.77±0.11 nu, p<0.001, Y: 0.87±0.11 nu, p<0.001).

**Fig. 6.**
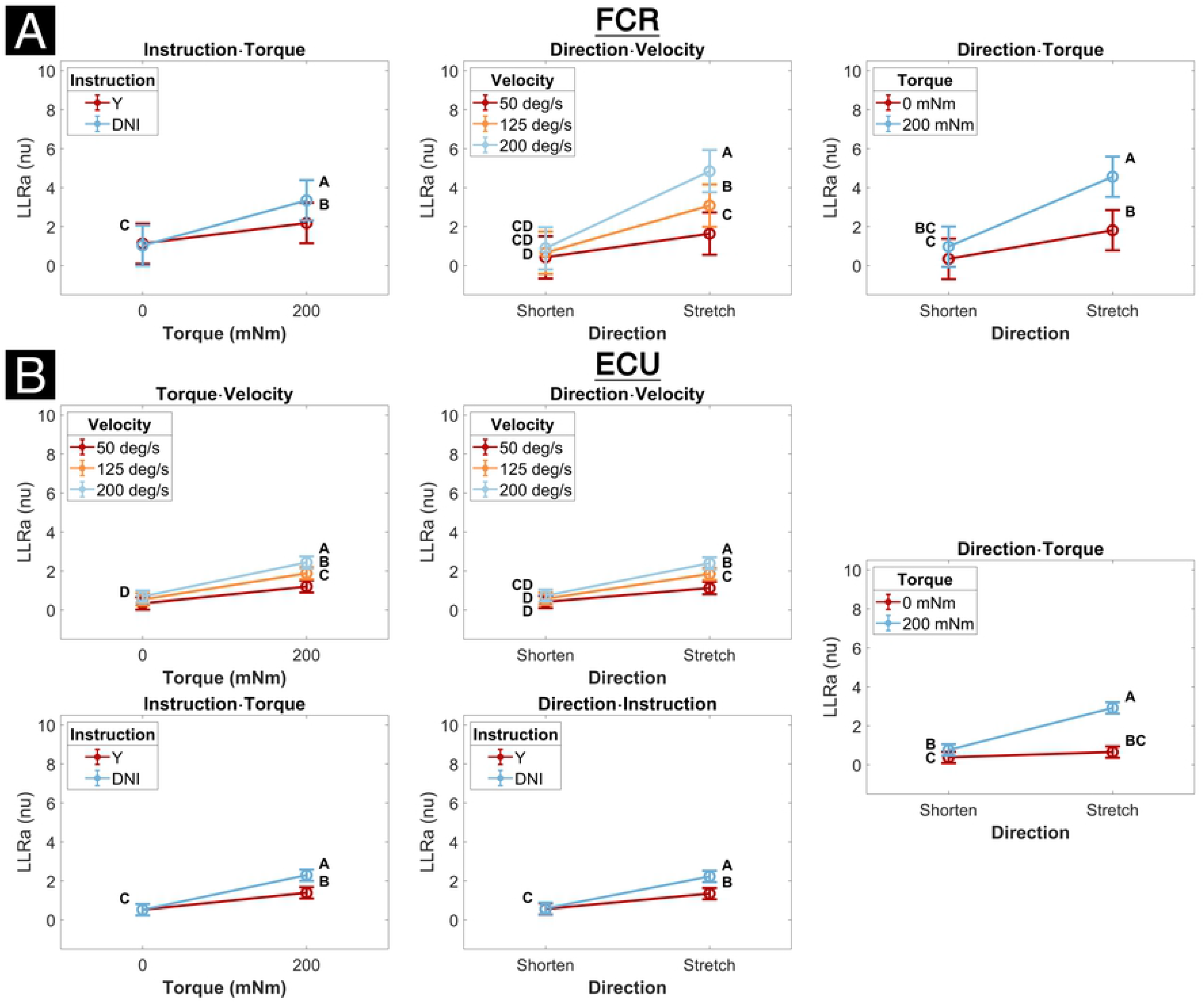
Least square means for all significant two-way interactions for both the FCR (top half) and the ECU (bottom half). Letters on the plots are representative post-hoc Tukey HSD tests; pairs of elements that do not share a letter are significantly different. Post-hoc tests are and the corresponding letter notation are separate for each panel.

The two-way interaction between perturbation direction and velocity resulted from a greater increase in LLRa measured when muscles are stretched by a perturbation compared to when they are shortened measured at higher velocities (Fig. 6). LLRa measured during stretch were significantly greater at higher velocities (FCR LLRa — 200deg/s: 5.31±0.56 nu, 125 deg/s: 3.37±0.56 nu, 50 deg/s 1.79±0.56 nu, 200−125 deg/s p<0.001, 125−50 deg/s p=0.003; ECU LLRa — 200 deg/s: 2.40±0.16 nu, 125 deg/s 1.85±0.16 nu, 50 deg/s 1.13±0.16 nu, 200−125 deg/s p=0.002, 125−50 deg/s p<0.001), while there was no significant effect of velocity during perturbations where the muscle was shortened. The increase in LLRa between shorten and stretch conditions was greater at higher velocities (FCR ΔLLRa — 50 deg/s: 1.34±0.42 nu, 125 deg/s 2.66±0.42 nu, 200 deg/s 4.35±0.42 nu, 200−125 deg/s p=0.005, 200−50 deg/s p<0.001, 125−50 deg/s p=0.026; ECU ΔLLRa — 50 deg/s: 0.71±0.14 nu, 125 deg/s 1.27±0.14 nu; 200 deg/s 1.66±0.14 nu, 200−125 deg/s p=0.047, 200−50 deg/s p<0.001, 125−50 deg/s p=0.006).

The significant two-way interaction between perturbation direction and torque results from the greater increase in LLRa associated with stretch perturbations compared to shortening measured in presence of background torque (Fig. 6). LLRa associated with stretch perturbations were greater than those associated with shortening in presence of background torque (FCR ΔLLRa — 0 mNm: 1.62±0.34 nu, 200 mNm: 3.95±0.34 nu, p<0.001; ΔLLRa ECU — 0 mNm: 0.27±0.11 nu, 200 mNm: 2.15±0.11 nu, p<0.001). LLRa measured in presence of background torque were greater than in absence of background torque for both muscles when they were stretched (FCR ΔLLRa stretch: 3.02±0.34 nu, p<0.001; ECU ΔLLRa stretch: 2.26±0.11 nu, p<0.001), but only for ECU when shortened (FCR ΔLLRa shorten: 0.69±0.34 nu, p=0.180; ECU ΔLLRa shorten: 0.38±0.11 nu, p=0.006).

Two two-way interaction terms were significant only for the ECU muscle. One term, the interaction between torque and velocity, resulted from a greater increase in LLRa associated with higher velocity perturbations measured in the presence of a background torque (Fig. 6). The change in LLRa associated with the two levels of background torque increased at greater velocities (ECU ΔLLRa — 200 deg/s: 1.75±0.14 nu, 125 deg/s: 1.34±0.14 nu, 50 deg/s: 0.87±0.14 nu, 200−125 deg/s p=0.040, 200−50 deg/s p<0.001, 125−50 deg/s p=0.0181). LLRa measured in presence of background torque was significantly different for each velocity level and increased with greater velocity (ECU ΔLLRa — 200−50 deg/s: 1.24±0.14 nu, p<0.001, 200−125 deg/s: 0.56±0.14 nu p=0.001, 125−50 deg/s: 0.68±0.14 nu p<0.001).

The second two-way interaction term significant only the for ECU muscle was the two-way interaction between perturbation direction and instruction. This term resulted from a greater increase in LLRa measured in the DNI condition compared to yield condition measured when muscles are stretched (Fig. 6). When muscles were stretched, LLRa associated with the DNI conditions were larger than the yield condition (ECU LLRa — DNI: 2.23±0.15 nu, Y: 1.35±0.15 nu, p=0.001), while no significant difference between instruction conditions was measured when muscles were shortened. A significant increase in LLRa was measured in both instruction conditions when muscles were stretched compared to shortened, but this increase was greater in the DNI condition compared to the yield condition (ECU ΔLLRa — DNI: 1.64±0.11 nu; Y: 0.79±0.11 nu, p<0.001).

The model returned a significant effect of each of the four factors, i.e. perturbation direction, instruction, background torque, and velocity. For the effect of perturbation direction, stretched muscles resulted in a significantly larger LLR amplitude than those of shortened muscles (FCR LLRa — stretch: 3.49±0.50 nu, shorten: 0.71±0.50 nu, p<0.001; ECU LLRa — stretch: 1.79±0.13 nu, shorten: 0.58±0.13 nu, p<0.001). With respect to the significant effect of instruction, the DNI condition resulted in larger LLR amplitudes compared to the yield condition for both the FCR and ECU (FCR LLRa — DNI: 2.39±0.50 nu, yield: 1.81±0.50 nu, p=0.016; ECU LLRa — DNI: 1.41±0.13 nu, yield: 0.96±0.13 nu, p<0.001). The presence of background torque at 200 mNm resulted in a significantly larger LLR amplitude compared to the absence of background torque for both muscles (FCR LLRa — 200 mNm: 3.03±0.50 nu, 0 mNm: 1.17±0.50 nu, p<0.001; LLRa ECU — 200 mNm: 1.84±0.13 nu, 0 mNm: 0.52±0.13 nu, p<0.001). The significant effect of velocity resulted from higher velocities associated with larger LLRas for both muscles (FCR LLRa — 50 deg/s: 1.12±0.52 nu, 125 deg/s: 2.04±0.52 nu, 200 deg/s: 3.14±0.52 nu; ECU LLRa — 50 deg/s: 0.77±0.14 nu, 125 deg/s: 1.21±0.14 nu, 200 deg/s: 1.56±0.14 nu). Tukey HSD post-hoc analysis indicated that all velocity levels are significantly different from one another for both muscles. (FCR — 200−125 deg/s: p<0.001, 125−50 deg/s: p=0.006; ECU — 200−125 deg/s: p=0.002, 125−50 deg/s: p<0.001).

### 1D analysis

The results of the 1D ANOVA are represented in terms of a timeseries of F scores, shown in Fig. 7 and Fig. 8 for the ECU and FCR, respectively. For effects and interactions that have significant upcrossings within the LLR region, post-hoc 1D t-tests were used to further break down the effects.

**Fig. 7.**
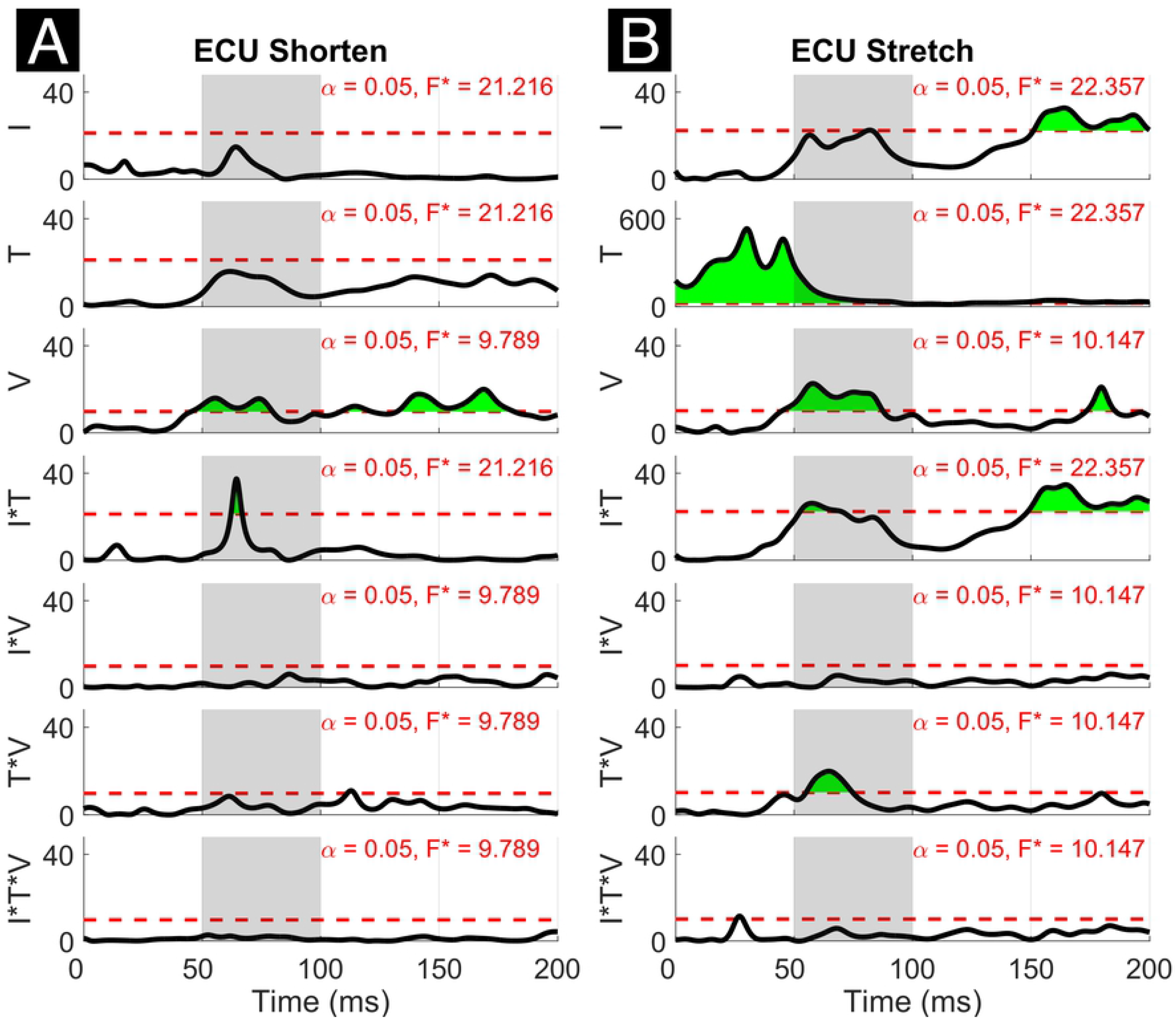
Results of the 1D 3-way ANOVA for the shortened (left) and stretched (right) states of the ECU. Curves are a time-series of F scores for the effect of a factor or of an interaction of factors on the outcome measure. The dashed red line indicates the critical F value for that main effect or interaction. Regions in green indicate significant upcrossings. The shaded gray region is representative of the LLR (50 to 100 ms).

**Fig. 8.**
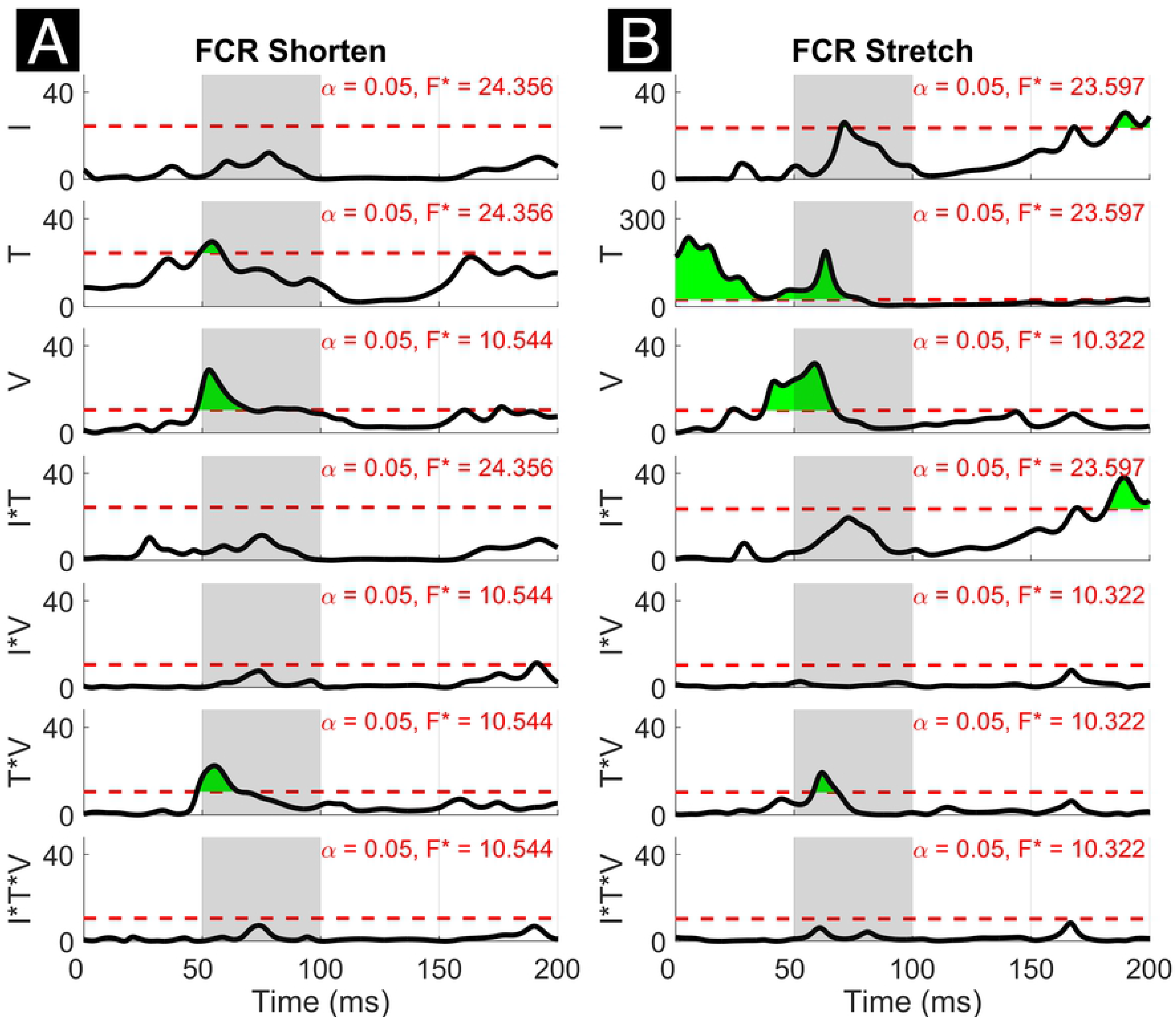
Results of the 1D 3-way ANOVA for the shortened (left) and stretched (right) states of the FCR.

The three-way interaction between instruction, torque, and velocity was not significant within the LLR for both muscles and both directions, however, there is a narrow significant upcrossing during the SLR for ECU Stretch (25.8 to 28.3 ms, peak F_2,20_ = 11.495).

The two-way interaction between instruction and torque is significant within the LLR region for both directions of the ECU muscle, but not for the FCR muscle. Within the LLR region, there are significant upcrossings for both the shortened and stretched states of the ECU (ECU Stretch: 52.6 to 70.0 ms, local peak in LLR region F_1,10_ = 26.152; ECU Shorten: 62.2 to 66.8 ms, peak F_1,10_ = 37.434). There are also significant upcrossing within the voluntary region for the stretch of the ECU and FCR (ECU Stretch: 148.6 to 200 ms, peak F_1,10_ = 26.152; FCR Stretch: 183.0 to 196.0 ms, 199.3 to 200 ms, peak F_1,10_ = 27.377).

The two-way interaction between instruction and torque measured for the ECU muscle is broken down in the 1D post-hoc t-tests to analyze the measured effect at all time points, as shown in Fig. 9. Analysis of post-hoc tests highlights how the DNI has positive or negative effect on the amplitude of processed EMG recordings at different time points, and the effect differs as a function of stretch and background torque condition. When the ECU was stretched, the normalized EMG signal in DNI was greater than yield within the LLR and voluntary regions when in the presence of background torque, (ECU Stretch: 49.1 to 200 ms, peak T = 13.703, Fig. 9B, center). No significant change in EMG signal was measured in absence of background torque as a function of task instructions (Fig. 9B, left). As a result, the change in EMG signal measured between the DNI and Y condition was greater in presence of background torque than in absence only after the delay similar to that considered for forearm LLRs (ECU Stretch: 38.7 ms to 200 ms, peak ΔT = 14.450, peak in the LLR region ΔT = 9.397, Fig. 9B, right). When the ECU was shortened, EMG recordings measured in presence of background torque in the DNI condition were greater than those measured in the yield condition in the initial part of LLR, but then EMG recordings measured during DNI were smaller than yield at a later time in the LLR time period (ECU Shorten: 57.7 to 77.0 ms, local peak T = 4.470, 94.6 ms to 129.5 ms, local peak T = –3.925, Fig. 9A, center). Instead, no significant change in EMG signal was measured in absence of background torque as a function of task instructions (Fig. 9A, left). The change in T scores between torque levels for the shortened condition also indicated significant upcrossings in the LLR and voluntary regions (ECU Shorten: 54.2 to 73.5 ms, local peak ΔT = 4.961, 92.5 to 132.9 ms, local peak T = −5.461, Fig. 9A, right).

**Fig. 9.**
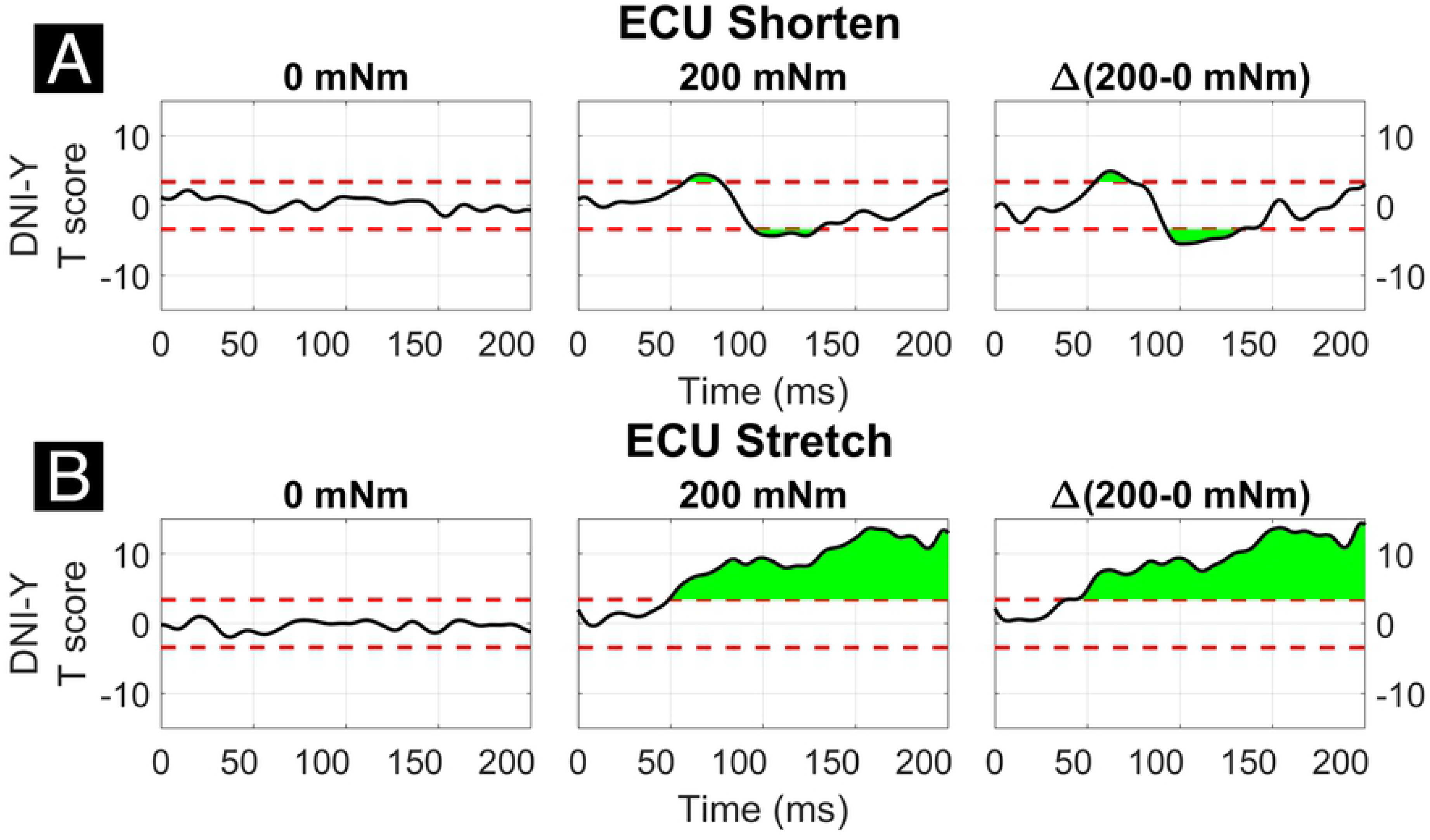
Results of the 1D t-tests for the interaction between task instruction and torque for the shortened (top) and stretched (bottom) states of the ECU. Plots indicate the statistical score of 1D paired t-tests between the DNI and yield instructions and split in columns by torque level. Plots in the right column report the difference between the T-scores in the second and first column’s tests, used to demonstrate interaction between these factors. The dashed red line indicates the Bonferroni-adjusted critical T values and regions in green indicate significant upcrossings.

The two-way interaction between torque and velocity is significant in the LLR region for the ECU when stretched and for the FCR when it is both shortened and stretched. For both the ECU and FCR there is a significant upcrossing within the LLR region (ECU Stretch: 52.6 to 74.7 ms, peak F_2,20_ = 19.872; FCR Shorten: 47.9 to 65.8 ms, peak F_2,20_ = 22.421; FCR Stretch: 58.0 to 68.6 ms, peak F_2,20_ = 19.244).

The two-way interaction between torque and velocity for muscle and perturbation direction is broken down in 1D post-hoc tests for ECU Stretch, FCR Shorten, and FCR Stretch (Fig. 10-12). For the stretch of the ECU (Fig. 10), significant upcrossings were present in post-hoc tests comparing EMG signals measured at different velocity levels and at multiple torque levels, primarily during the LLR time period. Analysis of the timeseries of t-scores demonstrate that EMG signal increases with velocity and with background torque, and that the region where such an increase is measured largely overlap with the expected latency of a LLR. The largest upcrossing was measured for the 200-50 deg/s comparison in presence of a background torque (40.8 to 98.1 ms, peak T = 10.202, Fig. 10, center row, center column). Upcrossings within the LLR region were also measured for the 200-50 deg/s t-test in the absence of background torque, the 200-125 deg/s t-test in the presence of background torque, and the 125-50 degree/second t-test in both torque conditions. The significance of the between-torque condition difference of the t-scores resulting from comparing pairs of velocity levels indicates that background torque differentially modulates the velocity dependence of EMG signals. Significant upcrossings were measured within the LLR region for the 200-50 deg/s comparison and the 200-125 degrees/second comparison (Δ200-50 deg/s: 56.4 to 73.8 ms, peak ΔT = 5.915, Fig. 10 center row, right column, Δ200-125 deg/s: 56.0 to 62.7 ms, peak ΔT = 4.304, Fig. 10, top row, right column).

**Fig. 10.**
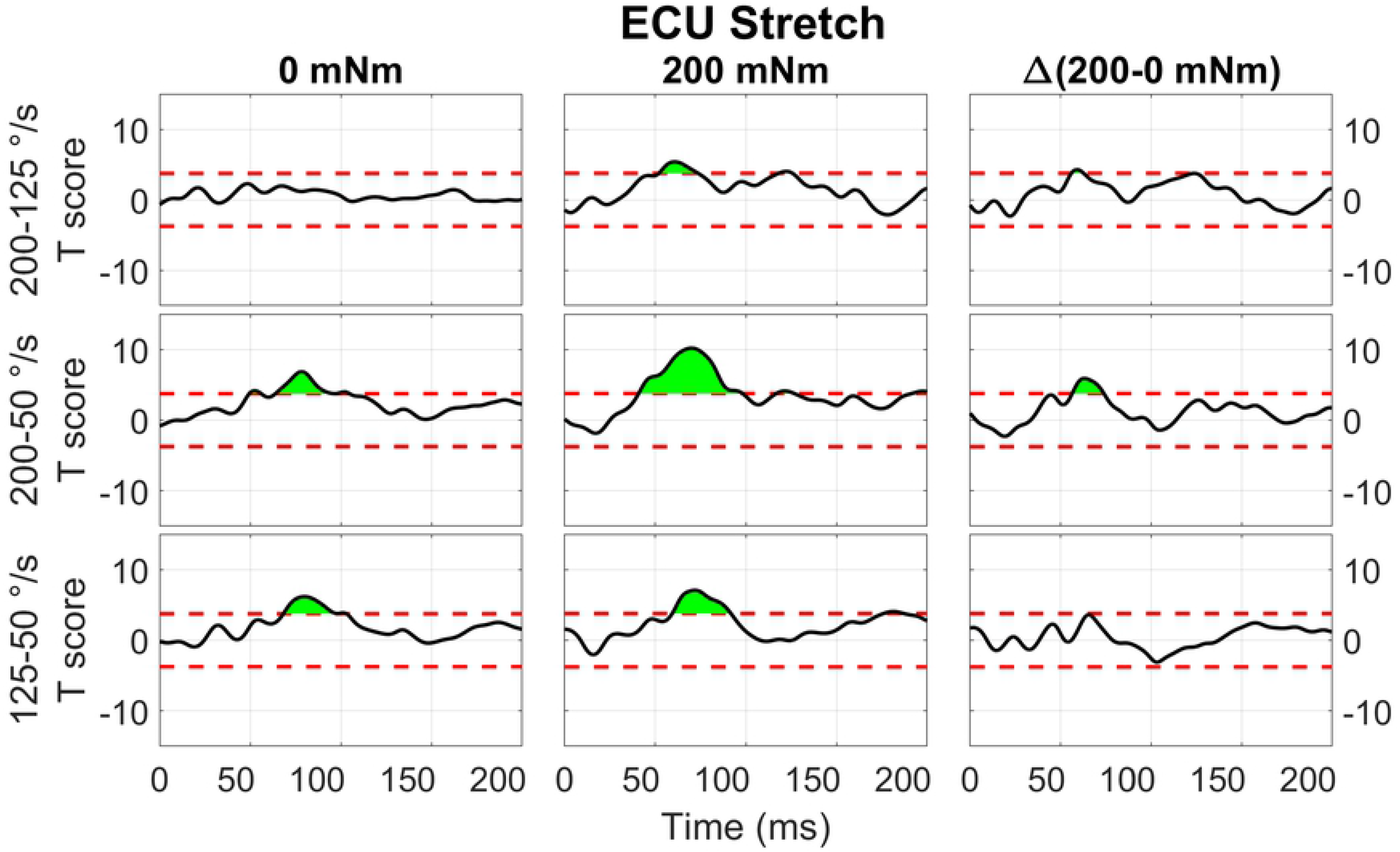
Results of the 1D t-tests of the interaction between torque and velocity for the stretched state of the ECU. Plots in different rows indicate the statistical score of 1d paired t-tests between the velocity levels (200-125, 200-50, 125-50 deg/s) and columns indicate different torque levels. Plots in the right column report the difference between the T-scores in the second and first column’s tests, used to demonstrate interaction between these factors. The dashed red line indicates the Bonferroni-adjusted critical T values and regions in green indicate significant upcrossings.

Qualitatively similar results were measured for the stretch of FCR (Fig. 11). Significant upcrossings were measured for t-test comparisons mostly during the LLR time period. The largest t-scores were generated for the 200-50 deg/s t-test in the presence of background torque (23.7 to 80.9 ms, peak T = 12.856, Fig. 11 center row, center column). Significant upcrossings were also present for the 200-125 deg/s t-test in presence of background torque, 200-50 degree/second t-test in the absence of background torque, the 200-125 deg/s t-test in both torque conditions, and the 125-50 deg/s t-test in both torque conditions. One significant upcrossing was measured for the between-torque condition difference of t-scores resulting from comparing pairs of velocity conditions (Δ200-50 deg/s: 57.9 to 64.0 ms, peak ΔT = 4.604, Fig. 11 center row, right column).

**Fig. 11.**
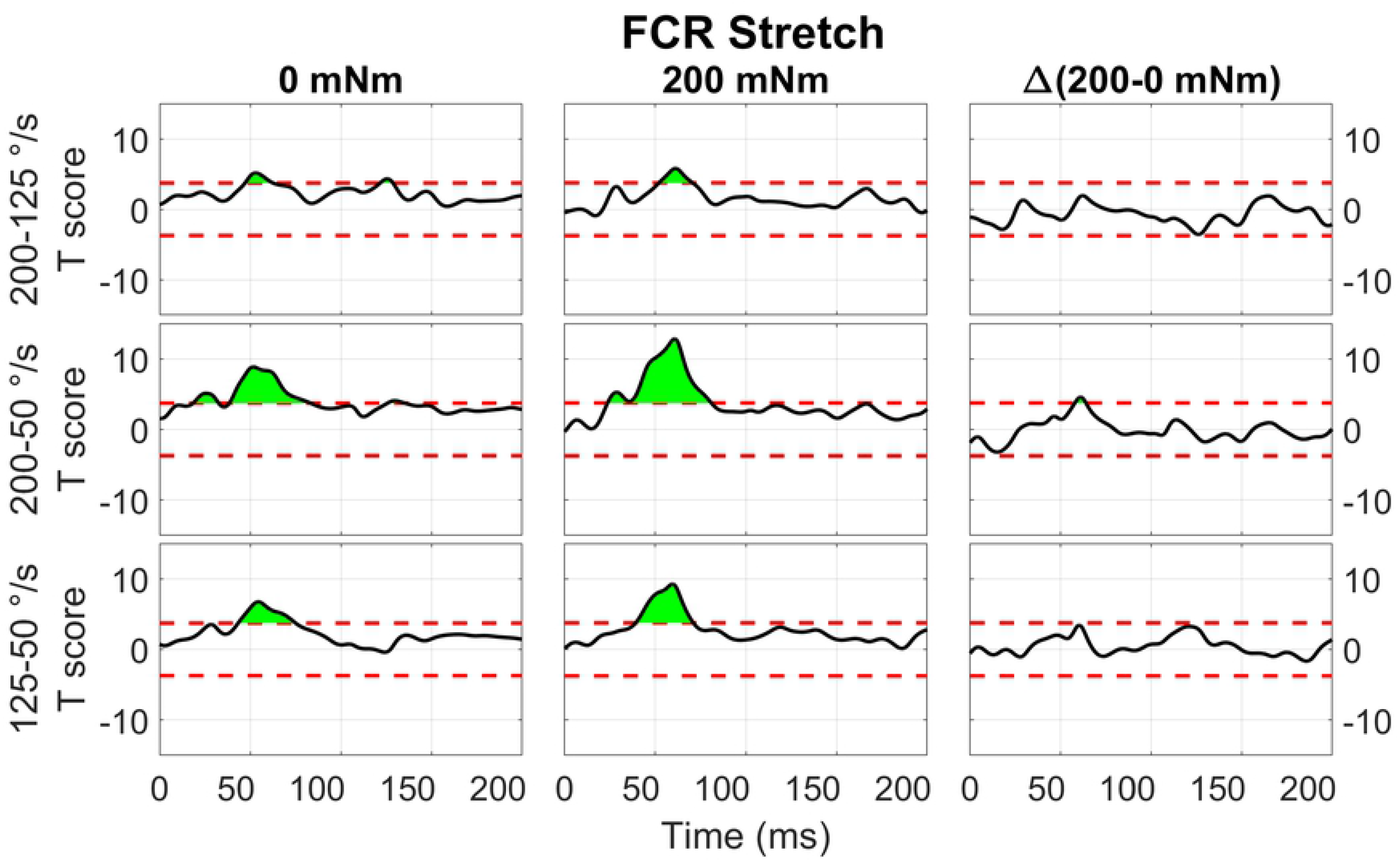
Results of the 1D t-tests of the torque and velocity interaction for the stretched state of the FCR

With respect to the shortened state of the FCR (Fig. 12), significant upcrossings were present in t-test comparisons between velocity levels in the presence of background torque. Like in the previous conditions, the largest t-scores were generated for the 200-50 degree/second t-test in the presence of background torque (46.0 to 93.3 ms, peak T = 8.601, Fig. 12 center row, center column). There are also smaller, significant upcrossing for this t-test in the voluntary region (149.1 to 165.1 ms, 187.9 to 200 ms). Significant upcrossings were also present for the 200-125 deg/s t-test in presence of background torque, 200-50 degree/second t-test in the absence of background torque, and the 125-50 degree/second t-test in both torque conditions. Significant upcrossings within the LLR region were calculated for difference between t-tests of different background torque levels (Δ200-50 deg/s: 48.7 to 63.2 ms, peak ΔT = 6.587, Fig. 11 center row, right column, Δ200-125 deg/s: 50.6 to 59.2 ms, peak ΔT = 5.176, Fig. 11, top row, right column). Such differential effect of background torque at multiple velocities on EMG signal collected from a muscle under shortening was only observed for the ECU muscle in the 0D analysis.

**Fig. 12.**
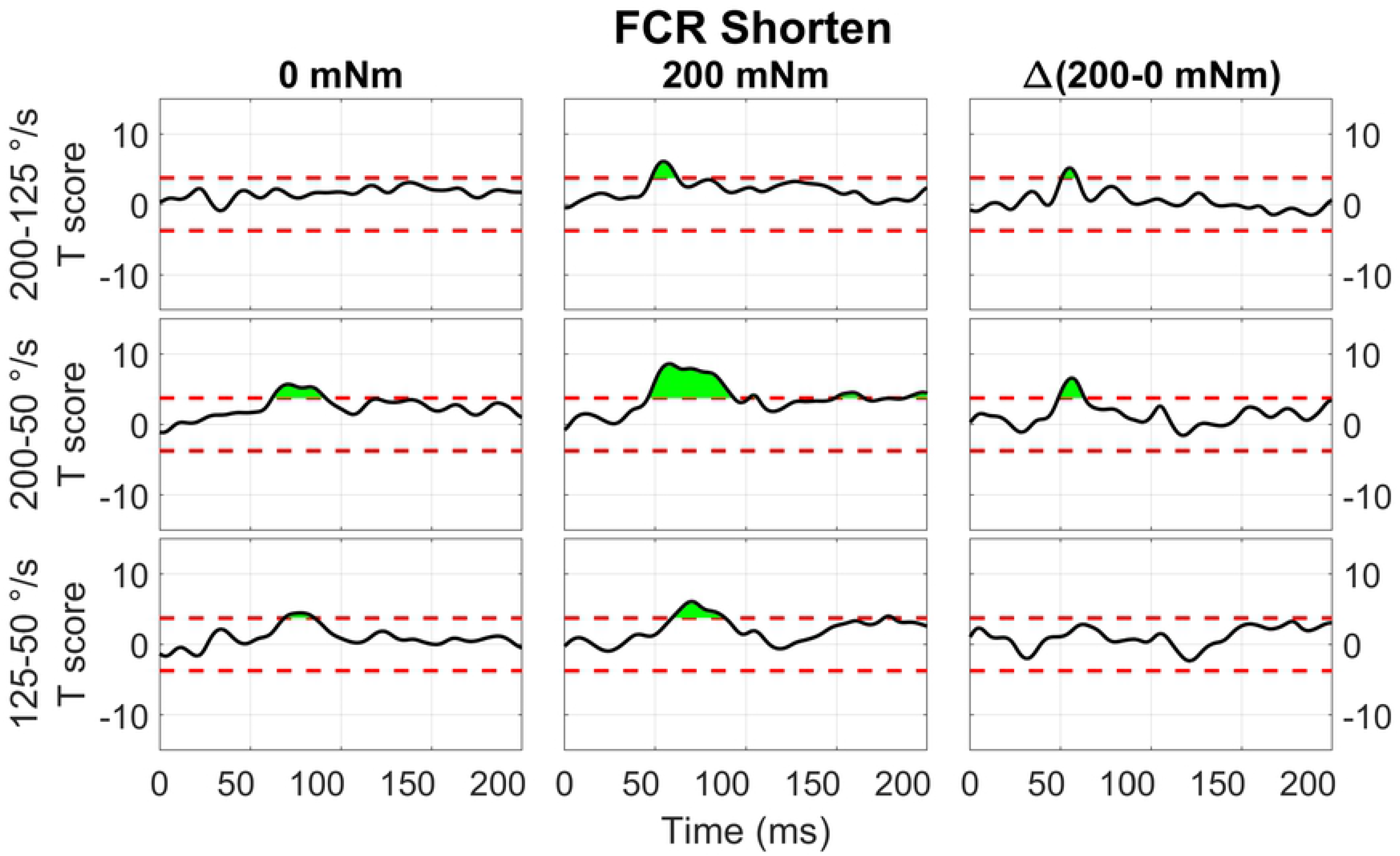
Results of the 1D t-tests of the torque and velocity interaction for the shortened state of the FCR.

The two-way interaction between instruction and velocity is only significant for the shortened state of the FCR during a narrow time window outside of the LLR region (189.8 to 193.1 ms, peak F_2,20_ = 10.959).

With respect to the main effects, both the FCR and ECU had significant upcrossings within the LLR region. The main effect of instruction was significant within the LLR region for the stretch of the FCR and ECU only for a very short time period (FCR stretch: 69.8 to 73.2 ms, local peak F_1,10_ = 26.080; ECU stretch: 81.1 to 82.8 ms, local peak F_1,10_ = 22.546) and has upcrossings within the voluntary region for both the stretch of the FCR and ECU (FCR Stretch: 184.6 to 200 ms, peak F_1,10_ = 30.620; ECU Stretch: 151.1 to 200 ms, peak F_1,10_ = 32.771). There were otherwise no significant upcrossings for the effect of instruction for the shortened state of the muscles.

## Discussion

In the present study, we aimed to investigate the effects of four behavioral factors in modulating the LLR amplitude (LLRa) evoked from FCR and ECU muscles during ramp-and-hold perturbations applied to the wrist joint. The four behavioral factors studied in this work were perturbation direction (stretch vs. shorten), background muscle activation (0 vs. 200 mNm of joint torque requiring agonist muscle activity), perturbation velocity (50 deg/s, 125 deg/s, 200 deg/s), and task instructions (yield vs resist). In line with previous studies (15–24), we observed that all factors modulate the LLRa in the upper limbs’ muscles. However, given the use of full factorial design involving all combinations of the four factors, we were able to study and observe higher-order interactions between the four behavioral factors subject to investigation that had not been established previously. Moreover, the use of a 1D analysis of stretch-evoked EMG data allowed us to establish the effects of the behavioral factors on EMG amplitude without an a-priori hypothesis on when the modulation could be measured.

In line with previous studies (15,27), our findings demonstrated that perturbation velocity significantly modulates LLRa, as larger LLRa resulted from perturbations with 200 deg/s velocity compared to the 125 and 50 deg/s velocity conditions, while the 50 deg/s perturbation velocity evoked the lowest LLRa. Note that given the chosen task design, where perturbations are applied for a constant duration (200 ms), perturbation velocity and displacement are associated, so the measured effect could be due to the fact that the joint is subject to a greater displacement in the higher velocity conditions. From studies that systematically spanned a set of perturbation velocity and durations (20–22), we know that total joint displacement is the primary contributor to LLRs, with perturbations at high velocity, but brief duration not generating significant LLRa. In our experience, the execution of LLRa studies on perturbation durations that require stopping the limb within the [50 100] ms interval proved to be challenging, as in preliminary data we observed the occurrence of motion artifacts associated with the oscillations induced by the manipulator stopping. For this reason, we decided to focus on perturbations of longer duration (200 ms), which allowed to completely avoid those artifacts. Results also revealed a significant increase in LLRa when a 200 mNm background torque was applied prior to the perturbations in comparison to the perturbations with no background torque. Pre-existing background muscle activation is thought to reflect an automatic adjustment mechanism, known as the automatic gain component of LLR (15,19,25,26). Task instruction also significantly modulated the LLR in this study. The majority of previous studies examined the ‘yield’ or ‘DNI’ instruction with a ‘resist’ or ‘compensate’ instruction (19,23,24); However, to the best of our knowledge, except the study by Calancie et al. in 1985 (16), no prior study compared if ‘yield’ and ‘DNI’ task instruction would modulate the LLRa. Our findings showed that larger LLRs were evoked when participants attempt keep constant the muscles’ activation state (i.e., DNI), compared to when they are asked to yield to the perturbations, which is justifiable in accordance to the evidence that the temporal overlap of two different responses including a task-dependent response and an automatic response results in the task-dependent change in LLR amplitude (13,23,30). In line with a few previous studies quantifying LLR from both stretched and shortened muscles (18,19), the above-mentioned modulations had more prominent effects on the LLRa evoked from the muscle stretched in response to the perturbations vs. the shortened muscle.

With our full factorial design, we were able to observe more granular modulations of the LLR response induced by the four behavioral factors reported above. Specifically, significant results were observed for all the three-way interaction terms that can be computed without including the interaction between instruction and velocity, as detailed below.

### Perturbation direction, task instruction and background torque

As illustrated in Fig. 4, perturbation direction did not affect the LLRa evoked from the perturbations with no background torque. This finding is in contrast with a previous study by Miscio et al. (19), which demonstrated that there would be a lower LLRa evoked from the shortened muscle in comparison to the LLRa evoked from the stretched muscle even in absence of background muscle activity. In addition, task instruction did not modulate the LLRa evoked in absence of background torque, as in this case the behavior elicited in the Y and DNI conditions is expected to be identical.

Results from the perturbations with 200mNm background torque showed a significant difference between LLRa evoked from stretched muscle in the Y and DNI task instructions. Larger LLRa was detected from perturbations when participants were instructed to maintain their level of activation (DNI) after the perturbation, compared to when they were instructed to yield to the displacement. In contrast, task instructions did not modulate the LLRa evoked from the shortened muscle neither in presence of a 200mNm background torque. Also, the LLRa for shortened muscle was significantly lower than the LLRa for stretched muscle in both instruction conditions. This interaction is also consistently reported also in the 1D analysis (Fig. 9).

In agreement with this result, Miscio et al. (19) also reported a smaller LLRa for shortened muscle in comparison to the stretched muscles when there was a 10% maximum voluntary contraction (MVC) background torque and subjects were instructed to not intervene to the perturbations. However, their task instructions did not include a ‘yield’ condition for comparison. The EMG activity recorded from the shortened muscle in the previous study was interpreted as a volume conducted response based on their direct nerve stimulation results. Besides, their regression analysis showed a positive correlation between the area of FCR stretch response and the low-level activity recorded from the shortened antagonist muscle (ECR) (r2= 0.59). Instead, our data supports a differential modulation of the response for the stretched and shortened muscles between different levels of task instructions, suggesting that the source of the observations may not be due to cross-talk, but to true physiological decoupling of the muscle responses.

### Perturbation direction, background torque, perturbation velocity

Results of the 0D analysis (Fig. 5) support a significant interaction between the effects of torque and velocity on the LLRa evoked from perturbations stretching the ECU muscle, but not from perturbations shortening the ECU muscle. In support of this observation, the 1D analysis (Fig. 10) revealed a significant interaction between torque and velocity on the LLRa for ECU muscle only under stretching, but not under shortening. For the FCR muscle, the interaction was significant in response to the both stretching and shortening perturbations, though the differential effects of background activity are smaller, and last for a smaller extent for this muscle (Fig. 11, 12). The smaller size and duration of the effects of torque and velocity, combined with the greater between-subject variability of FCR data might be a possible reason why the 0D analysis did not detect a significant effect of the torque and velocity interaction for the FCR muscle.

Previous studies also demonstrated a significant effect of the interaction between perturbation torque and velocity on the LLRa evoked from the stretched muscle (15). However, based on our literature review, no prior study evaluated this interaction for the a muscle subject to a perturbation that shortened the muscle. Berardelli et al. (34), reported that background torque and velocity would increase the amplitude of the EMG activity evoked from the shortened tibialis anterior muscle with a latency within 100-150ms, but no interaction analysis was conducted in that study.

For both muscles, perturbation direction did not modulate the LLRa elicited by any velocity condition in absence of the background torque. However, Miscio et al. (19), observed lower LLRa for the shortened muscle compared with the stretched muscle which were perturbed with 500 degree/s velocity with no background torque. It might be possible that higher perturbation velocities are needed to differentiate the long latency stretch response of the agonist and antagonist muscles when there is no background activity. Moreover, to the best of our knowledge, our study is the first study compared the effects of different perturbation velocities on the LLRa between the shortened and stretched upper limbs’ muscles, and the first to observe an interaction between background torque and velocity in the LLRa of a muscle undergoing shortening, an effect that was visible for the FCR muscle. Therefore, further studies are needed to make a strong justification on this topic.

### Interaction between task instruction and velocity

No analysis has revealed any interaction between the task instruction and perturbation velocity affected the LLRa. The wajority of previous work studied the interaction between perturbation velocity with perturbation duration and amplitude (20–23). Among them, Lewis et al. (23), was the only study that examined the interaction between the task instruction and perturbation velocity. In contrast to our results, they reported a significant interaction (p=0.02), which resulted in a greater facilitation in the LLRa evoked from the Biceps brachii muscle, for the ‘Flex’ instructed perturbations with a 90 degree/s velocity. In their ‘Flex’ instruction, subjects were instructed to flex their elbows as soon as possible in response to the perturbation, a condition which wasn’t included in our experiment. Hence, it is possible that countering perturbations applied to the proximal upper limb joints (such as the elbow) would elicit a different modulation in the LLR from the one observed here for forearm muscles.

Our finding of no interaction between task instructions and perturbation velocity might be a direct consequence of the fact that distinct neural pathways are involved in producing the response associated with velocity and task instructions. In fact, there is a similar velocity-sensitive response in the SLR, which involves a monosynaptic pathway from the muscle spindle group Ia afferents to the motoneuron (21). Additionally, there is a body of evidence that the LLR evoked from the distal upper limb muscles receive more inputs from the transcortical loop compared to the proximal muscles (35,36), which can also explain the difference between the velocity and instruction interaction results for Lewis et al. study on elbow and our findings for the wrist muscles. Although the velocity-sensitive response is cortically-mediated, studies demonstrated that task instruction does not modulate the EMG activity over the SLR time period, which support the idea that modulation of the task-dependent component of LLR derived down stream from the motor cortex and involve a greater contribution from the subcortical motor regions as the brainstem (3,37).

### Similarity of the effects between FCR and ECU

Some of the significant interaction terms were only significant for ECU, and not for FCR. Specifically, no three-way interaction term was significant for the FCR muscle (as opposed to two terms significant for FCR), and two of the five two-way interactions significant for the ECU muscle were not significant for FCR. Overall, the mismatch of results between the two muscles is primarily due to larger variability of the measurements obtained for the FCR muscle compared to the ECU muscle, while all effects were generally in the same direction for both muscles. In many cases, in fact, the magnitude of measured modulations in LLRa (expressed in normalized units, so roughly comparable between muscles) was actually larger for the FCR muscle than for the ECU muscle. However, the standard error of the mean estimated by the mixed model was several times larger for the FCR muscle compared to the ECU muscle.

As an example, the term corresponding to the three-way interaction between direction, instruction, and torque was close to the chosen false I error rate for the FCR muscle (p=0.055). Further inspection of the parameter estimates corresponding to that interaction term shows that the mean difference in LLRa measured between Y and DNI conditions is slightly larger in the FCR muscle than it is for the ECU muscle under stretch (FCR LLRa – DNI: 6.05±0.58 nu, Y: 3.95±0.58 nu; ECU LLRa – DNI: 3.80±0.17 nu, Y: 2.04±0.17 nu,), while smaller effects were measured in the shorten condition (FCR LLRa – DNI: 1.28±0.58 nu, Y: 0.83±0.58 nu; ECU LLRa – DNI: 0.78±0.17 nu, Y: 0.73±0.17 nu) for both muscles. Yet, the four-fold larger s.e.m. measured for the FCR muscle reduces the statistical power in detecting this effect.

Another interaction term was close to the conventional type I error rate in the 0D analysis for FCR, i.e., the interaction between perturbation direction and instruction (p=0.13). Once again, the difference in modulation between shortening and stretching perturbations at the two task instruction levels was larger in the FCR muscle compared to ECU (FCR – ΔLLRa DNI: 3.14±0.34 nu, Y: 2.42±0.34 NU; ECU – ΔLLRa DNI: 1.64±0.11 nu, Y: 0.79±0.11 nu). Yet, the larger variability of the slopes measured in the FCR was responsible for the smaller significance of the measured effect.

### 0D vs. 1D analysis

To the best of our knowledge, this study is the first to apply a mixed-model 1D analysis to the timeseries of EMG signals measured in response to velocity-controlled perturbations. 1D analyses are used to quantify if and when a timeseries is significantly modulated by an experimental factor, without a-priori hypotheses on the duration of this modulation, or on the time window where such a modulation is expected. Typically, reflex studies use 0D variables such as the average of the rectified signal in a pre-defined time window (29), or the cumulative sum of the rectified EMG signal within a region of supra-threshold response (31) – the latter definition allowing to explicitly account for a combined measure of amplitude and duration of a reflex response.

A shortcoming of using 0D datapoints generated from a 1D data set is the possibility for false positives. It has been indicated that the false positive rate of the 0D data is greater than the desired false positive rate of α = 0.05 when using noisy outcome measures such as peak in an interval (32), and that this may be the case also in presence of smoothing (32,33). Less noisy outcome measures such as the mean EMG signal in a pre-defined time window are not likely to result in limited control of the false positive rate of inference tests. However, the main advantage of using a 1D analysis for the study of stretch reflexes is the fact that the 1D analysis may give a more granular insight on the shape of the waveform (e.g. one peak, two peak) which can be masked in taking an average to create a 0D dataset, and/or to test hypothesis on whether an effect is ONLY present in a specific time-region, while controlling for the type-I error rate of inferential statements.

As an example, our 0D analysis showed that, in presence of background activation, there is an effect of task instruction on the LLRa measured from the ECU when stretched, but no effect measured when the muscle is shortened (Fig. 4). However, a closer inspection on this modulation is provided by the post-hoc t-tests presented in Fig. 9. It is true that the difference between rectified EMG signal measured in the DNI and Y condition for the ECU under stretch is greater in the 200 mNm condition, reflecting a larger signal measured in the DNI condition (Fig. 9, center). However, the same contrast plot extracted from the ECU during shortening perturbations highlights the presence of a biphasic response, where the DNI condition results in an early increase in EMG signal in the LLR time window, followed by a decrease in EMG signal. The average of the positive and negative changes is likely the reason why the same effect could not be captured using the 0D analysis.

Also, insight from a 1D analysis provides information about the gradient in timing resulting from different behavioral factors. Specifically, the effect of instruction is usually visible later than the one of velocity and torque, which have upcrossings in SLR and early LLR (Fig. 7 and Fig. 8). Instruction has a narrow region of significant main effects in the late LLR period (70 to 73 ms for FCR stretch, 81 to 83 ms for ECU stretch), with significant upcrossings in the voluntary period. Velocity conditions are significant in the early to mid LLR period (FCR stretch ~40 to 70 ms, ECU stretch ~45 to 90 ms, FCR shorten ~50 to 90 ms, ECU shorten ~45 to 80 ms), all having significant peaks near 60 ms. The 1D analysis allows us to deduct the presence of a 10 to 20 ms delay between the significant peaks due to instruction versus velocity. The time frame of these significant peaks are comparable with previous findings and the identification of two distinct peaks within the LLR period, named M2 and M3 in the literature. However, it is difficult to compare the exact timining in absolute terms since “latency” is often defined relative to the onset of EMG activity (16,19,22–24).

Yet, the results of the 0D and 1D analyses do not perfectly overlap. This is in part also due to limitations in the current version of the software used for the analysis. The spm1D MATLAB package allows up to three input conditions, as seen through its anova3rm function. To analyze the four conditions studied, two analyses per muscle were conducted when in the shortened and stretched conditions. As such, none of the three-way interactions that were significant in the 0D analysis could be studied using the 1D method. Another limitation of the 1D analysis is related to its limited statistical power. Because of the large number of datapoints included in the analysis, and the fact that the EMG signal is somewhat noisy and composed of multiple resolution elements in a time window of 200 ms following a perturbation, 1D analyses afford a reduced statistical power compared to a 0D analysis that only looks at the average within a pre-defined time window (33).

## Conclusions

In summary, this study has quantified the effect of four behavioral factors -- background torque, task instruction, perturbation velocity and direction -- on the long-latency response (LLR) amplitude evoked from the FCR and ECU muscles during ramp-and-hold perturbations applied to the wrist joint in the flexion and extension direction. Our analysis demonstrated that all of those factors modulate LLRa, and that their combination nonlinearly contribute to modulating the LLRa. Specifically, all the three-way interaction terms that can be computed without including the interaction between instruction and velocity significantly modulated the LLR. The interaction analysis suggested that higher background torque augmented the LLRa evoked from the stretched muscle when subjects are asked to maintain their muscle activation in response to the perturbations. Besides, higher perturbation velocity increased the LLRa evoked from the stretched muscle in presence of a background torque. While a lot of the behavioral factors listed above nonlinearly contribute to modulating LLRa, we observed that the effects of task instruction and velocity do not combine more than linearly to modulate the LLRa. Coherent with previous observations on the role of subcortical areas in modulating the task-dependent component of an LLRa, our results suggest that the modulation introduced by perturbation velocity and task instruction are processed by distinct neural pathways.

## Acknowledgement

Research reported in this publication was supported by the National Institute of Neurological Disorders and Stroke of the National Institutes of Health under award number R21NS111310. The content is solely the responsibility of the authors and does not necessarily represent the official views of the National Institutes of Health. Additional support was provided by the University of Delaware Research Foundation grant no. 16A01402, by ACCEL NIGMS IDeA grant no. U54-GM104941.

